# Mechanisms of distributed working memory in a large-scale network of macaque neocortex

**DOI:** 10.1101/760231

**Authors:** Jorge F. Mejias, Xiao-Jing Wang

## Abstract

To elucidate the circuit mechanism of persistent neural activity underlying working memory that is distributed across multiple brain regions, we developed an anatomically constrained computational model of large-scale macaque cortex. We found that persistent activity may emerge from inter-areal reverberation, even in a regime where none of the isolated areas is capable of generating persistent activity. The persistent activity pattern along the cortical hierarchy indicates a robust bifurcation in space, characterized by a few association areas near dynamical criticality. A host of spatially distinct attractor states is found, potentially subserving various internal processes. The model yields testable predictions including the idea of counterstream inhibitory bias, and suggests experiments to differentiate local versus distributed mechanisms. Simulated lesion or optogenetic inactivation revealed that distributed activity patterns are resilient while dependent on a structural core. This work provides a theoretical framework for identifying large-scale brain mechanisms and computational principles of distributed cognitive processes.

## Introduction

With the advances of brain connectomics and physiological recording technologies like Neuropixels^1,2^, a major new challenge in Neuroscience is to investigate biological mechanisms and computational principles of cognitive functions that engage many interacting brain regions. Here, the goal is no longer to identify one local parcellated brain region that contributes to or is crucial for a particular function, but how a large-scale brain system with many interacting parts underlies behavior. Currently, there is a dearth of theoretical ideas and established models for understanding distributed brain dynamics and function.

A basic cognitive function recently shown to involve multiple brain areas is working memory, the brain’s ability to retain and manipulate information in the absence of external inputs. Working memory has been traditionally associated with persistent neural firing in localized brain areas, such as those in the frontal cortex^3–8^ and computational models uncovered the involvement of local recurrent connections and NMDA receptors in the encoding of memory items via selective persistent activity^9–11^. However, self-sustained neural persistent activity during working memory has been found in multiple brain regions; and often such highly engaged areas appear in coactivation^12–14^. Previous modeling efforts have been limited to exploring either the emergence of persistent activity in local circuits, or in two interacting areas at most^15,16^. It is presently not known what biophysical mechanisms could underlie a distributed form of memory-related persistent activity in a large-scale cortex. The observation that mnemonic activity is commonly found in the prefrontal cortex (PFC) does not prove that it is produced locally rather than resulting from multi-regional interactions; conversely, a distributed persistent activity pattern could in principle be a manifestation of sustained inputs broadcasted by a local source area that can produce persistent activity in isolation. Therefore, understanding a distributed cognitive function is highly nontrivial and requires new theoretical ideas.

In this study, we tackle this challenge by building and analyzing an anatomically-constrained computational model of the cortical network of macaque monkey, and investigate a novel scenario in which long-range cortical interactions support distributed activity patterns during working memory. The anatomical data is used to constrain the model at the level of long-range connections but also at the level of local circuit connectivity. In particular, the model incorporated differences between individual cortical areas, by virtue of macroscopic gradients of local circuit properties in the large-scale network. The emerging distributed patterns of persistent activity involve many areas across the cortex. They engage temporal, frontal and parietal areas but not early sensory areas, in strong agreement with a recent meta-analysis of persistent activity in macaque cortex^13^. Persistent firing rates of cortical areas across the hierarchy display a gap, indicative of the existence of a robust bifurcation in cortical space. Furthermore, the distributed patterns emerge even when individual areas are unable to maintain stable representations, and increase the robustness of the network to distractors and simulated inactivation of areas. The concept departs from the classical view of working memory based on local attractors, and sheds new light into recent evidence on distributed activity during cognitive functions.

## Results

Our computational model includes 30 areas distributed across all four neocortical lobes (Fig. 1a; see Online Methods for further details). The inter-areal connectivity is based on quantitative connectomic data from tract-tracing studies of the macaque monkey^17–19^ (Extended Data Fig. 1). For simplicity, each of the cortical areas is modeled as a neural circuit which contains two selective excitatory populations and one inhibitory population^9,20^ (Fig. 1b). In addition, the model assumes a macroscopic gradient of synaptic excitation^21–23^ (Extended Data Fig. 2). This gradient is introduced by considering that the number of apical dendritic spines, loci of excitatory synapses, per pyramidal cells increases^24^ along the cortical hierarchy^18,25^ (Fig. 1c). This gradient of area-specific connection strength was applied to both local recurrent and long-range excitatory connections. In particular, we denote the maximal strength of local and long-range excitation for the area at the top of the cortical hierarchy by J_max_, which is an important parameter of the model (see below). To allow for the propagation of activity from sensory to association areas, we assumed that inter-areal long-distance connections target more strongly excitatory neurons for feedforward pathways and inhibitory neurons for feedback pathways, in a graded fashion^26^ (Fig. 1b). We shall refer to the gradual preferential targeting onto inhibitory neurons by top-down projections as the “counterstream inhibitory bias” hypothesis. It is worth noting that exploration of such new hypotheses would have not been possible without a quantitative definition of the cortical hierarchy and biologically-realistic circuit modeling.

**Figure 1:**
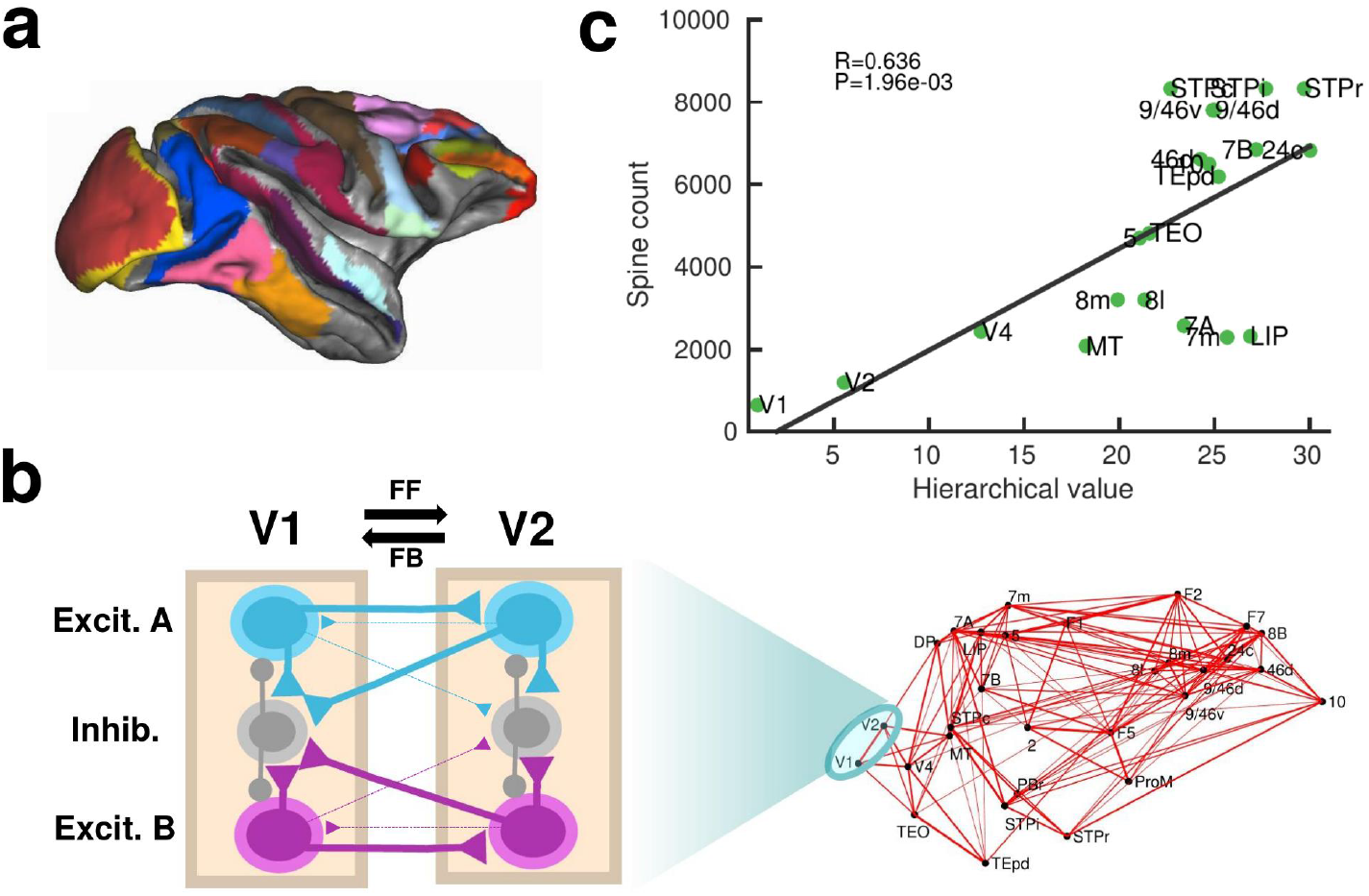
Scheme and anatomical basis of the multi-regional macaque neocortex model. (a) Lateral view of the macaque cortical surface with modelled areas in color. (b) In the model, inter-areal connections are calibrated by mesoscopic connectomic data^17^, each parcellated area is modeled by a population firing rate description with two selective excitatory neural pools and an inhibitory neural pool^27^. Recurrent excitation within each selective pool is not shown for the sake of clarity of the figure. (c) Correlation between spine count data^24^ and anatomical hierarchy as defined by layer-dependent connections^18^.

### Distributed WM is sustained by long-range cortical loops

In local circuit models of working memory (WM)^9,10^, areas high in the cortical hierarchy make use of sufficiently strong synaptic connections (notably involving NMDA receptors^9,11^) to generate self-sustained persistent activity. Specifically, the strength of local synaptic reverberation must exceed a threshold level (in our model, the local coupling parameter J_S_ must be larger than a critical value of 0.4655), for an isolated local area to produce stimulus-selective persistent activity states that coexist with a resting state of spontaneous activity (operating in a multistable regime rather than in a monostable regime, see Fig. 2a). However, there is presently no conclusive experimental demonstration that an isolated cortical area like dorsolateral prefrontal cortex (dlPFC) is indeed capable of generating mnemonic persistent activity. In this study, we first examined the scenario in which all areas, including dlPFC (9/46d) at the top of the hierarchy, have J_S_ values below the critical value for multistability (so *J*_*s*_ ≤ *J*_*max*_ < 0.4655) and are connected via excitatory long-range projections of global coupling strength G (we set J_max_ =0.42 and G=0.48 unless specified otherwise)(Fig. 2a). In this case, any observed persistent activity pattern must result from inter-areal connection loops. In a model simulation of a visual delayed response task, a transient visual input excites a selective neural pool in the primary visual cortex (V1), which yielded activation of other visual areas such as MT during stimulus presentation (Fig. 2b, upper left). After stimulus withdrawal, neural activity persists in multiple areas across frontal, temporal and parietal lobes (Fig. 2b, top right). The resulting activation pattern shows an excellent agreement with a large body of data, from decades of monkey neurophysiological experiments, reviewed in recent meta-analyses ^12,13^ regarding which areas display WM-related activity during delay period of WM tasks (Fig. 2b, bottom right). The activation pattern from the model was stimulus specific, so only the neural pool selective to the presented stimulus in each cortical area displayed elevated persistent activity (Fig. 2c; Extended Data Fig. 3). We observed cross-area variations of neural dynamics: while areas like 9/46d displayed a sharp binary jump of activity, areas like LIP exhibited a more gradual ramping activity, resembling temporal accumulation of information in decision-making^28^.

**Figure 2:**
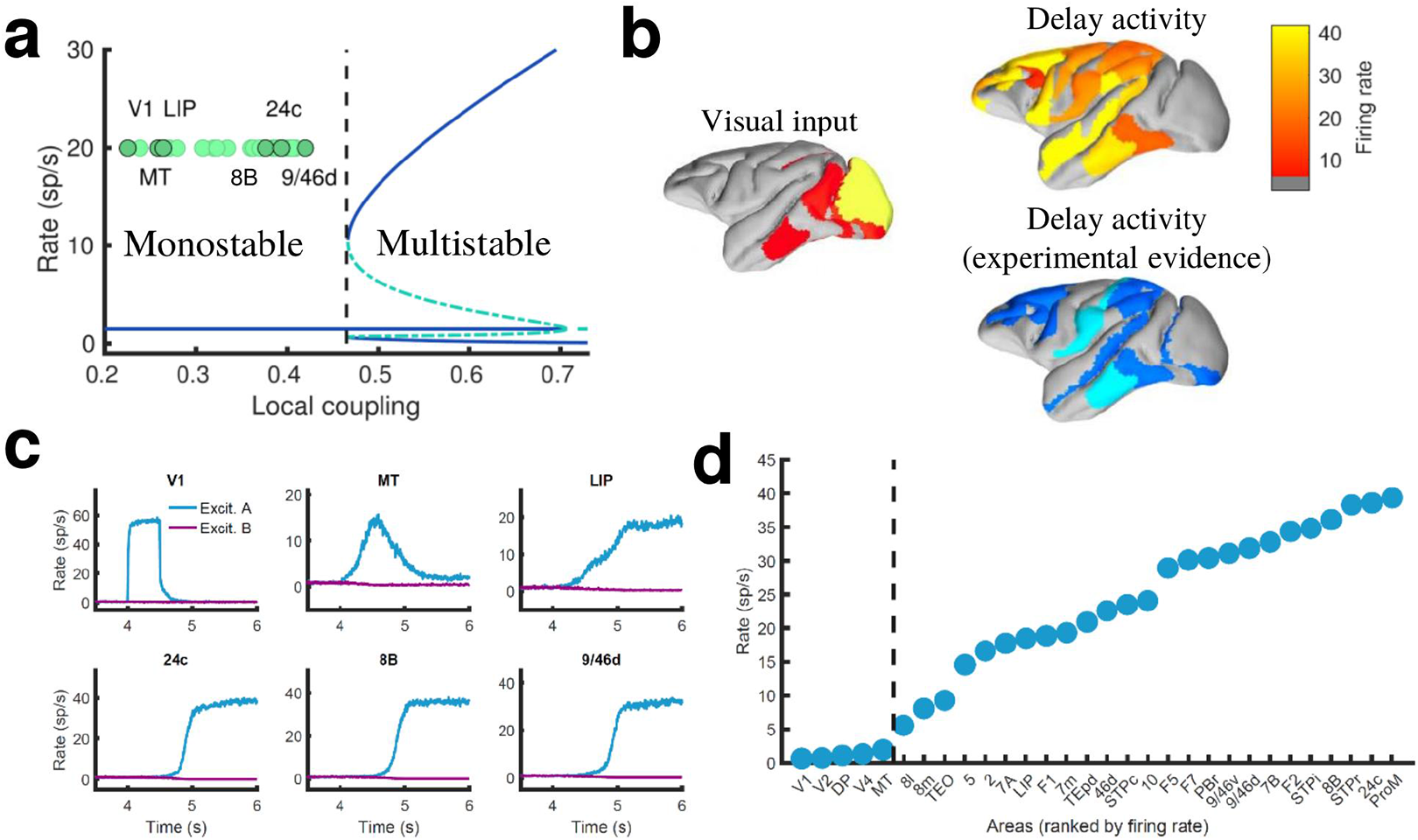
Distributed WM sustained via long-range loops in cortical networks. (a) Bifurcation diagram for an isolated area. Green circles denote the position of each area, with all of them in the monostable regime when isolated. (b) Spatial activity map during visual stimulation (left) and delay period (upper right). For comparison purposes, bottom right map summarizes the experimental evidence of WM-related delay activity across multiple studies^**13**^, dark blue corresponds to strong evidence and light blue to moderate evidence. (c) Activity of selected cortical areas during the WM task, with a selective visual input of 500 ms duration. (d) Firing rate for all areas during the delay period, ranked by firing rate.

Given that selective persistent activity is also found in somatosensory WM tasks^6^, we further test our model and simulate a simple somatosensory WM task by transiently and selectively stimulating a neural pool in primary somatosensory cortex. As in the case of visual stimulation, this leads to the emergence of a distributed persistent activity pattern of equal selectivity as the input (Extended Data Fig. 4), showing the validity of the distributed WM mechanism across different sensory modalities. Likewise, our model has considered NMDA receptors as the only excitatory dynamics for simplicity. However, AMPA dynamics may also be important^29^, and can be easily introduced leading to a good behavior of the model for shorter durations of the stimulus presentation (Extended Data Fig. 5).

When we plotted the firing rate of stimulus-selective persistent activity across 30 areas along the hierarchy, our results revealed a separation between the areas displaying persistent activity and those that did not (Fig. 2d). This is a novel type of bifurcation or transition of behavior that takes place in space, rather than as a function of a network parameter like in Fig. 2a. As a matter of fact, the relevant parameter here is the strength of synaptic excitation that varies across cortical space, in the form of a macroscopic gradient. The bifurcation is robust in two respects. First, the separation between areas appears not only when areas are ranked according to their firing rates, but also when they follow their positions in the anatomical hierarchy or in the rank of spine count values (Extended Data Fig. 6). Second, it does not depend on any fine tuning of parameter values.

### Simplified model of distributed working memory

The above model, albeit a simplification of real brain circuits, includes several biologically realistic features, which makes it difficult to identify essential ingredients for the emergence of distributed WM. For this reason, we developed a minimal model consisting on a fully connected network of excitatory firing-rate nodes (Fig. 3a, see Online Methods). The network includes a linear gradient of local properties: areas at the beginning of such gradient have weak self-coupling, while areas at the end have strong self-coupling. As in the more elaborated model, self-excitation is too weak to generate bistability in any isolated nodes.

**Figure 3:**
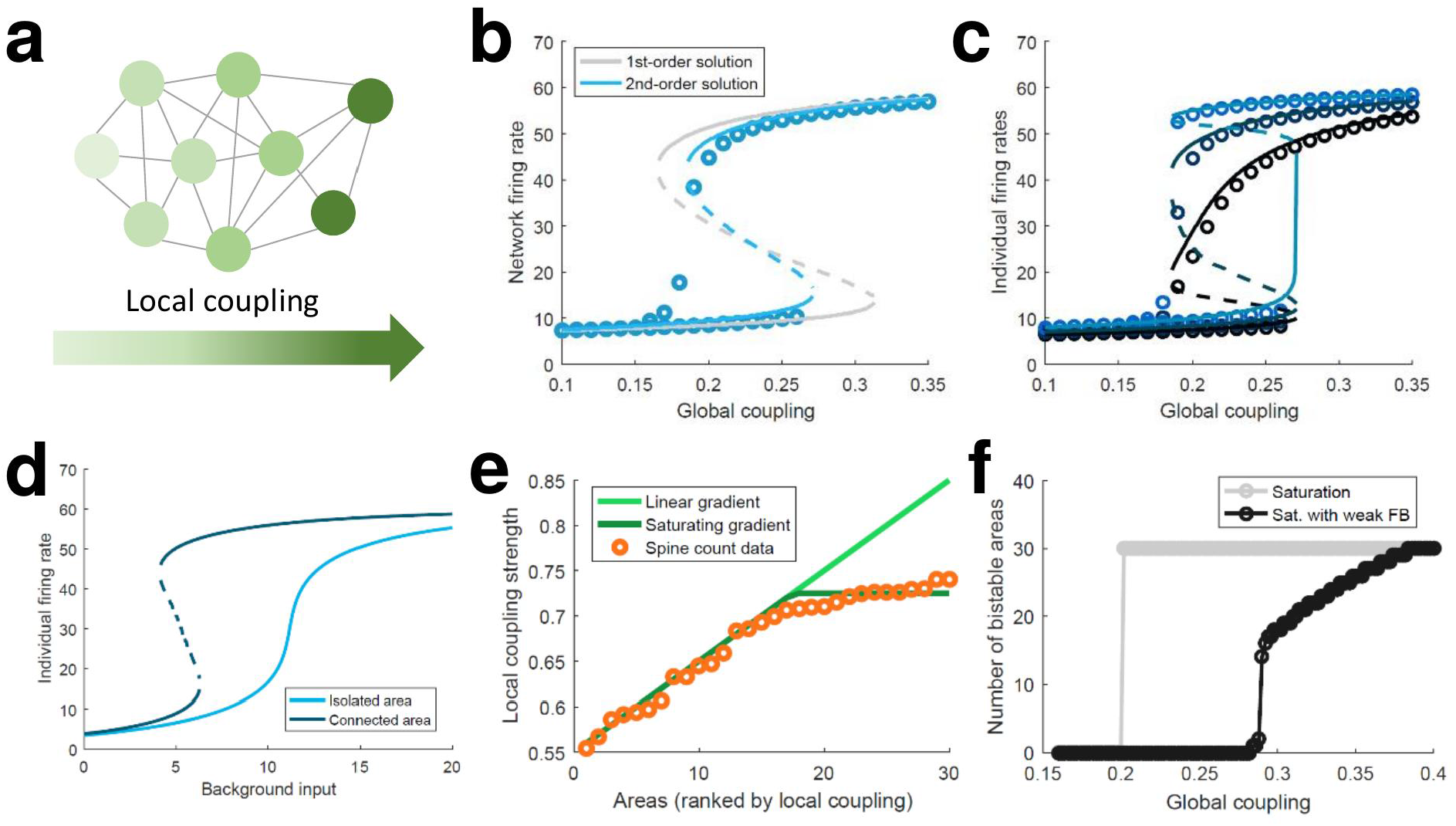
Simplified model of distributed WM. (a) Scheme of our simplified model: a fully connected network of N=30 excitatory nodes with a gradient of local coupling strengths. (b) Population-average firing rate as a function of the global coupling strength, according to numerical simulations (symbols) and a mean-field solution based on first-order (grey line) or second-order (blue) statistics. (c) Firing rates of three example individual nodes (symbols denote simulations, lines denote second-order mean-field solutions). (d) Activity of an example node when isolated (light blue) or connected to the network (dark blue). (e) Two forms for the gradient of local coupling strength (lines) compared with the spine count data (symbols). (f) The number of bistable areas in the attractor (grey) is either zero or N when the gradient of local properties saturates as suggested by spine count data. When feedback projections become weaker, resembling inhibitory feedback, the network is able to display distributed WM patterns which involve only a limited number of areas (black).

This simple model allows for a mean-field analytical solution for the network average firing rate R of the form: *R* = *ϕ* ((*Jη*_0_ + *G*) *R* + *I*), with *ϕ* being a sigmoidal function, *Jη*_0_ the average local coupling value across areas, *G* the inter-areal connection strength, and *I* a background input current (see Online Methods for a full derivation of a two-area example and a more complete N-area network). The factor *Jη*_0_ + *G* determines whether the above equation presents one stable solution (spontaneous firing) or two (spontaneous and persistent firing). As this factor includes both local and global components, the average network firing rate may be bistable even if local couplings are weak, as long as the inter-areal connections are strong enough. This mean-field solution, as well as a more precise second-order version, show a good agreement with numerical simulations and confirm the emergence of distributed activity in the system (Fig. 3b). Simulations also show, around *G*∼0.17, the appearance of states in which only areas at the top of the gradient show bistability, indicated by low values of R. Once R is known, the mean-field solution also permits to predict the emergence of persistent activity in individual nodes (Fig. 3c) and also observe how monostable isolated units become bistable when incorporated into the network (Fig. 3d).

The simplified model demonstrates that distributed WM patterns, in which some (but not all) areas display bistability when connected to each other, may emerge on generic networks of excitatory units as long as (i) their long-range connections are strong enough and (ii) the network has a linear gradient of local couplings. When considering biological constraints, however, these two conditions might not be easy to meet. In particular, data on the area-specific number of spines per neuron seems to monotonically increase, but saturates instead of growing linearly (Fig. 3e). Introducing this saturating gradient on the simplified model makes the nodes more homogeneous, and as a result the network is not able to display distributed WM patterns without indistinctively activating all nodes (Fig. 3f, grey curve). This problem was solved when we assumed that feedforward projections (i.e. those going from lower to higher areas in the gradient) were slightly stronger while feedback projections were slightly weaker, which is consistent with the counterstream inhibitory bias hypothesis. Such assumption, needed for saturating gradients, allows to recover solutions in which only a subset of areas display bistability in the WM patterns (Fig. 3f, black curve).

### Impact of the counterstream inhibitory bias in the full model

As indicated by the simplified model, introducing differences between feedforward and feedback projections is a key ingredient to achieve realistic patterns of distributed WM in a data-constrained model. In the full, biologically more realistic model, this asymmetry is introduced by considering a graded preferential targeting to inhibitory neurons by top-down projections (i.e. counterstream inhibitory bias, or CIB), which prevents indiscriminate persistent activation across all cortical areas^13^ (Fig. 4a). We systematically varied the strength of the feedback projections targeting inhibitory population in our model, and computed the firing rates of different areas during delay for these cases. We observed that, for strong enough CIB, the overall firing rate of early sensory areas is reduced, while the activity levels of areas high in the hierarchy is maintained at appropriate values (Fig. 4b). This also allows distributed WM patterns to emerge for a wide range of the global coupling strength (Fig. 4c).

**Figure 4:**
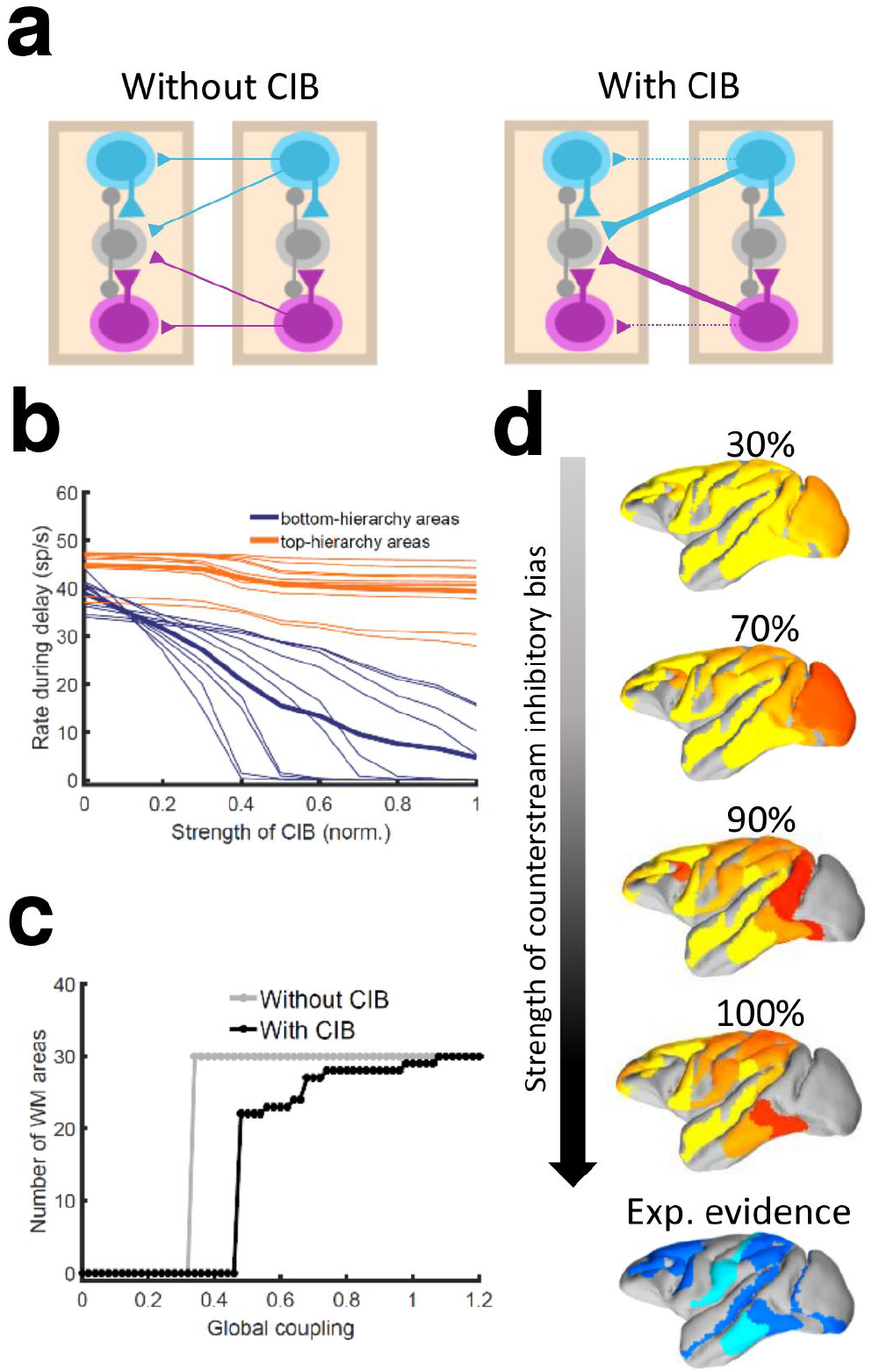
Effect of inhibitory feedback on distributed WM. (a) Scheme showing a circuit without (left) or with (right) the assumption of counterstream inhibitory bias, or CIB. (b) Firing rate of areas at the bottom and top of the hierarchy (10 areas each, thick lines denote averages) as a function of the CIB strength. (c) Number of areas showing persistent activity in the example distributed activity pattern vs global coupling strength without (grey) and with (black) CIB. (d) Activity maps as a function of the CIB strength. As in Fig. 2b, bottom map denotes the experimental evidence for each area (dark blue denotes strong evidence, light blue denotes moderate evidence).

The strength of the counterstream inhibitory bias has also an impact on the overall activity profiles across the brain network. Fig. 4d shows activity maps for several CIB levels, revealing that a moderate to strong bias shows a good agreement with the experimental evidence.

It is also worth noting that exceptions to the CIB rule may exist in brain networks without compromising the stability of distributed WM attractors. For example, a more balanced targeting would allow for WM-related activity in primary visual areas, which still constitutes a point of controversy in the field^13^. FEF areas 8l and 8m, on the other hand, are not able to sustain persistent activity when receiving strong inhibitory feedback (especially from other frontal areas) and had to be excluded from this general rule, although such exception does not affect the results aside from local effects in FEF (Online Methods, Extended Data Fig. 7).

In Fig. 2 and also in the following sections, the strength of the counterstream inhibitory bias was considered proportional to the fraction of infragranular projections, as suggested by anatomical studies^19^ and following previous work^26^. This results in a very small bias for most of the projections, but enough to produce the desired effect (see Online Methods for further details). In addition to supporting the emergence of distributed WM, CIB could explain observed top-down inhibitory control effects^30^.

While the macroscopic gradient of excitability is an important property of the model, the particular values of excitatory strength assigned to each area are not relevant for the phenomenon (Extended Data Fig. 8a-d). Similar conclusions can be obtained when the anatomical structure of the cortical network is changed, for example by randomly shuffling individual projection strength values (Extended Data Fig. 8e-h). However, in this case the duration of the persistent activity for multiple areas may be affected. This suggests that salient statistical features of the structure embedded in the cortical network may play a role in the emergence of distributed activity patterns. The model also predicts the emergence of a hierarchy of time scales across cortical areas (Extended Data Fig. 9), in agreement with experimental findings^31^ and supporting and improving previous computational descriptions^21^.

### Long-range cortical loops support a large number of different distributed attractors

We realized that a large-scale circuit can potentially display a large number of distributed persistent activity patterns (attractors), and some of them may not be accessible by stimulation of a primary sensory area. Note that distinct attractor states are defined here in terms of their spatial patterns, which does not depend on the number of selective excitatory neural pools per area. We developed a numerical approach to identify and count distinct attractors (see Online Methods for further details). Our aim is not to exhaustively identify all possible attractors, as the activity space is too large, but to gain insight on how our estimations depend on relevant parameters such as the global coupling strength G, or the maximum area-specific synaptic strength J_max_. Five examples of different distributed WM attractors are shown in Fig. 5a, where we can appreciate that not all distributed attractors engage cortical areas at all lobes, and that frontal areas are the ones more commonly involved.

**Figure 5:**
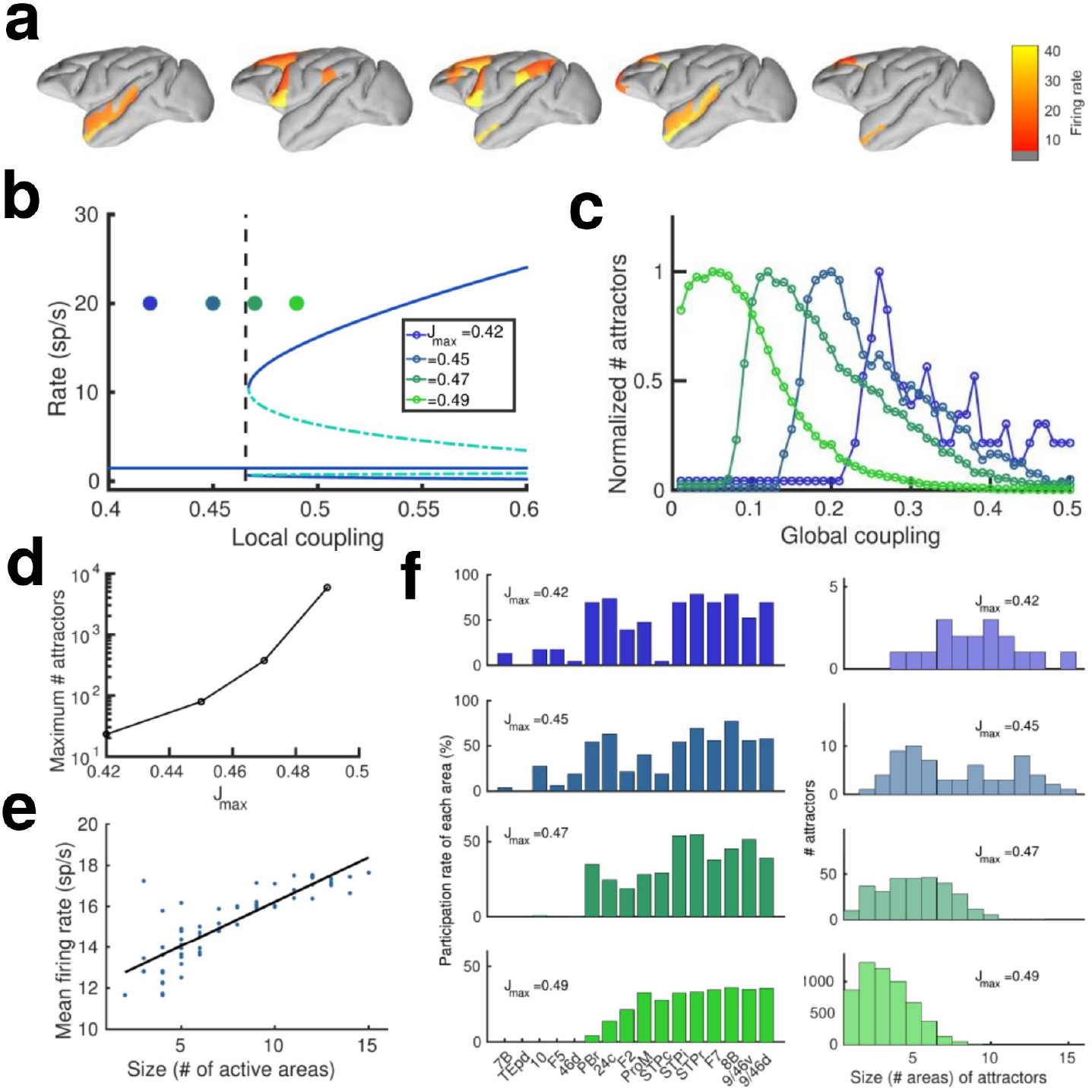
Distributed and local WM mechanisms can coexist in the model. (a) Five example distributed attractors of the network model. (b) Bifurcation diagram of an isolated local area with the four cases considered. (c) Number of attractors (normalized) found via numerical exploration as a function of the global coupling for all four cases. (d) Maximum (peak) number of attractors for each one of the cases. (e) Correlation between size of attractors and mean firing rate of its constituting areas for J_max_=0.45 and G=0.2. (f) Participation index of each area (left, arranged by spine count) and distribution of attractors according to their size (right).

A more detailed analysis included four cases depending on the value of the maximum area-specific synaptic strength J_max_ assumed: two of the cases had J_max_ above the bifurcation threshold for isolated areas (0.4655), and the other two had J_max_ below the bifurcation threshold. For the first two cases, having J_max_>0.4655 means that at least certain areas high in the hierarchy, such as dlPFC, have strong enough local reverberation to sustain activity independently (i.e. they were ‘intrinsically multistable’ and able to display bistability even when isolated from the network, Fig. 5b), however, areas lower in the hierarchy like 24c and F2 would require long-range support to participate in WM. For the last two cases, in which J_max_<0.4655, none of the areas was able to display bistability when isolated, but they can contribute to stabilize distributed WM attractors as in Fig. 2. In all four cases, the number of attractors turns out to be an inverted-U function of the global coupling strength G, with an optimal G value maximizing the number of attractors (Fig. 5c, curves are normalized to have a peak height of one for visualization purposes). This reflects the fact that a minimal global coupling is needed for areas to coordinate and form distributed WM attractors, but for values of G too large, all areas will follow the majority rule and the diversity and number of possible attractors will decrease. The optimal G value shifted towards lower values for increasing J_max_, and the peak number of attractors simultaneously increasing (Fig. 5d).

Across all four cases and G values considered, we found a significant positive correlation between the number of areas involved in a given attractor and the average firing rate of these areas (Fig. 5e), which constitutes an experimentally testable prediction of the distributed model of WM. We also analyzed how distributed WM attractors were constituted for the four different cases (Fig. 5f). When the network has a high number of intrinsically multistable areas (i.e. when J_max_>0.4655), attractors tend to only involve these areas and are therefore largely restricted to the areas located at the top of the hierarchy (Fig. 5f, bottom left and right panels). On the other hand, when the network has zero or a low number of intrinsically multistable areas (i.e. J_max_<0.4655), attractors typically involve a larger number of areas (as a larger pool of areas is needed to sustain distributed WM attractors, see top right panel in Fig. 5f) and the areas involved are more diverse in their composition (Fig. 5f, top left panel).

### Effects of inactivating areas on distributed attractors

To continue probing the robustness of distributed WM patterns, we tested the effect of inactivating cortical areas in our model during WM tasks, which can be done experimentally using optogenetic methods or lesioning selected areas. We tested this by completely and permanently suppressing the firing rate of the inactivated areas in the model, in such a way that the area becomes a sink of current and does not communicate with other areas. We began by inactivating (or silencing) a given number of randomly selected areas in a visually evoked distributed WM attractor, and found that the number of active areas in the attractor decreases only linearly with the number of inactivated areas (Fig. 6a). Furthermore, the activity of the areas remained in the distributed WM patterns linearly decreased their persistent activity level with the number of inactivated areas (Fig. 6b). When instead of selecting inactivated areas at random we silence areas in reverse hierarchical order (i.e. by inactivating top hierarchical areas first), the number of active areas decreases a bit more abruptly (Fig. 6c) and, as we will see later, can prevent the emergence of distributed WM altogether if J_max_ is sufficiently small.

**Figure 6:**
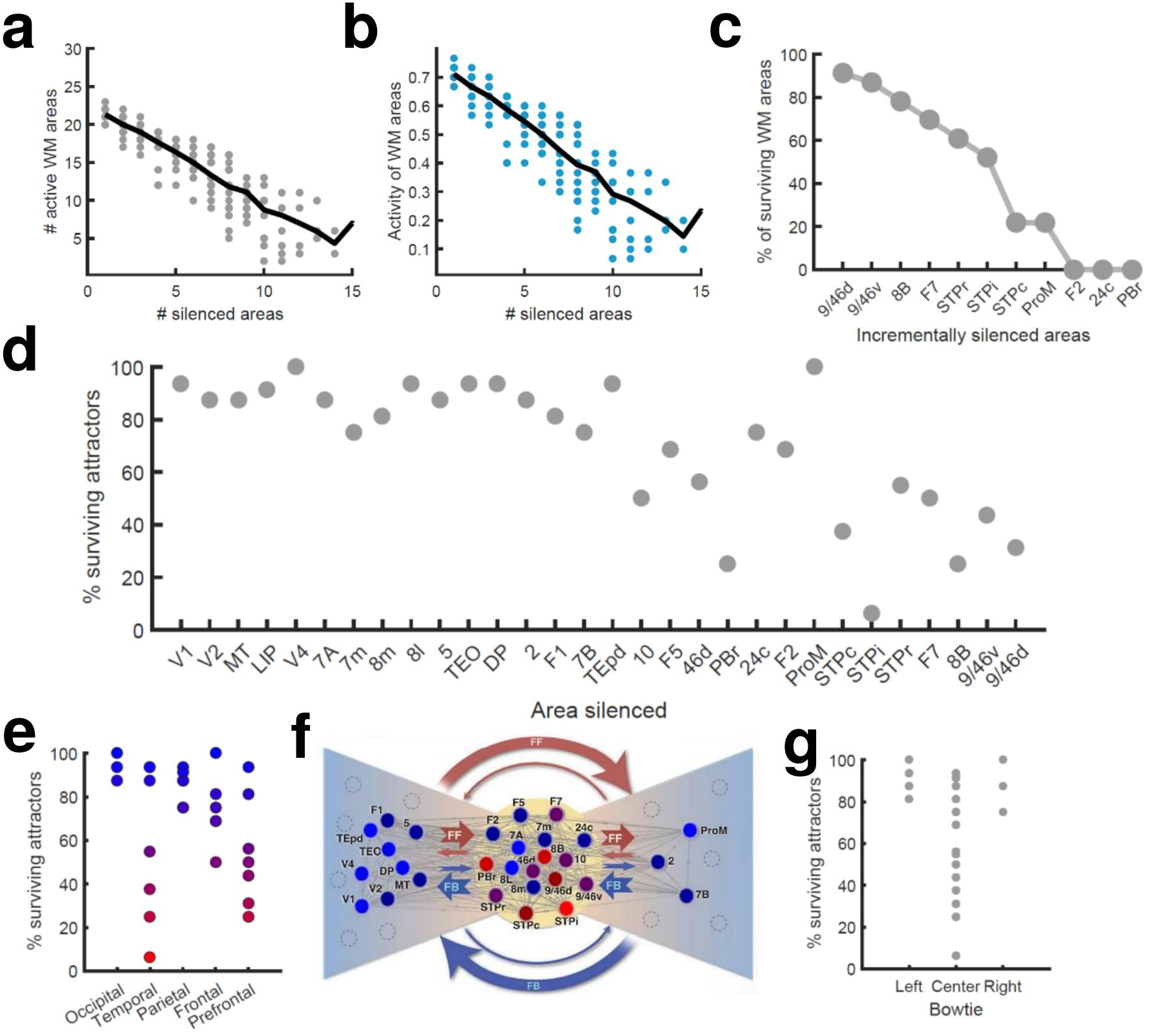
Effects of lesioning/silencing areas on the activity and number of attractors. (a) Number of active areas in the example attractor as a function of the number of (randomly selected) silenced areas. (b) The activity of the areas which remain as part of the attractor decreases with the number of silenced areas. (c) The number of active WM areas decreases faster when areas are incrementally and simultaneously silenced in reverse hierarchical order. (d) When considering all accessible attractors for a given network (G=0.48, J_max_=0.42), silencing areas at the top of the hierarchy has a higher impact on the number of surviving attractors than silencing bottom or middle areas. (e) Numerical exploration of the percentage of surviving attractors for silencing areas in different lobes. (f) Silencing areas at the center of the ‘bowtie hub’ has a strong impact on WM (adapted from^**17**^). (g) Numerical impact of silencing areas in the center and sides of the bowtie on the number of surviving attractors. For panels (e) and (f), areas color-coded in blue/red have the least/most impact when silenced, respectively.

We also carried out a more systematic evaluation of the effect of cortical inactivation, including their effect on attractors that were not accessible from sensory stimulation directly. This study revealed that inactivating most areas has only limited consequences on the total number of available distributed attractors, although in general the impact increases with the location of the silenced area in the hierarchy (Fig. 6d). In particular, the overall impact was large when some temporal and prefrontal areas are silenced, and sometimes more than half of the initially available attractors were lost (Fig. 6e). Interestingly, and beyond any hierarchical dependence, the temporal and prefrontal areas that had the strongest impact are part of the anatomical ‘bowtie hub’ of the macaque cortex identified in anatomical studies^17^ (Fig. 6f). Overall, silencing areas at the center of the bowtie had a deeper impact than silencing areas on the sides (Fig. 6g).

### Effects of area inactivation and distractors in distributed versus localized WM patterns

Across all analyses performed above, we assumed a relatively large value for the maximum area-specific recurrent strength J_max_=0.42, still below the critical value needed for bistability in isolation (0.4655). In order to provide clean predictions linked to the distributed WM scenario, in the following sections we studied the case of a strongly distributed WM system with J_max_=0.26 and G=0.48, and compared it to the case of networks which rely purely on a localized WM strategy (with J_max_=0.468, G=0.21 and feedback projections removed to avoid long-range loops).

We first reexamined the effect of inactivation for this strongly distributed WM network. We found that inactivation have in general a stronger effect here than for networks with larger J_max_ (as in Fig. 6). For example, inactivating key prefrontal areas such as 9/46d (dlPFC) fully prevented the emergence of distributed WM patterns evoked by external stimulation (Fig. 7a, b), which is in agreement with classical prefrontal lesion studies^32^. On the other hand, other areas can still be inactivated without disrupting distributed WM. In some cases, inactivating specific areas might even lead to a disinhibition of other areas and to a general reinforcement of the attractor (e.g., inactivating 24c leads to a larger and faster response by area STPi, Fig. 7b).

**Figure 7:**
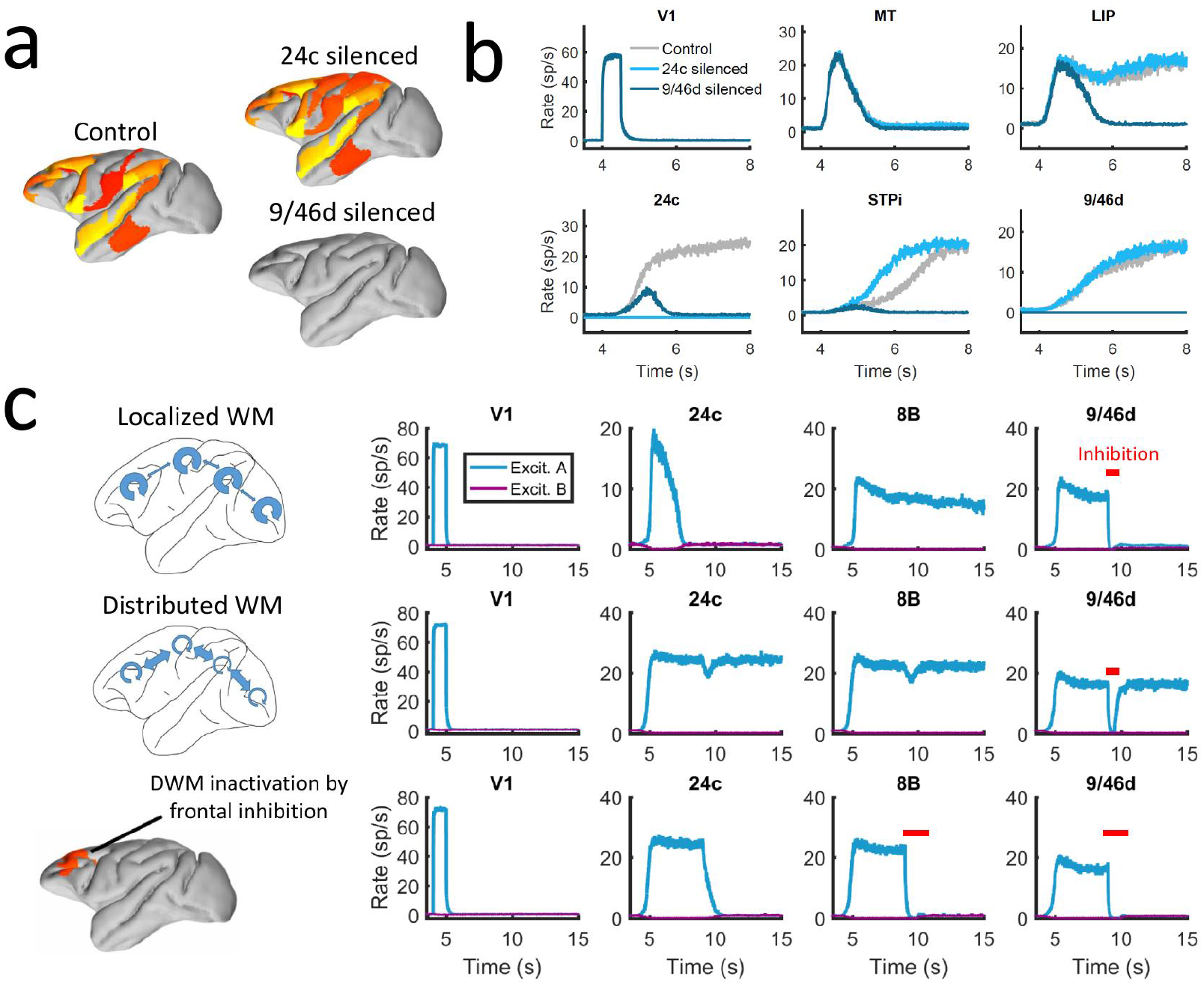
Effect of silencing areas in localized vs distributed WM. (a) Full-brain activity maps during the delay period for the control case (left), and lesioning/silencing area 24 (top right) or area 9/46d (bottom right). (b) Traces of selected areas for the three cases in (a) show the effects of silencing each area. (c) For a network displaying localized WM (top row, corresponding to J_max_=0.468, G=0.21), a brief inactivation of area 9/46d leads to losing the selective information retained in that area. For a network displaying distributed working memory (middle row, J_max_=0.26, G=0.48) a brief inactivation removes the selective information only transiently, and once the external inhibition is removed the selective information is recovered. In spite of this robustness to brief inactivation, distributed WM patters can be shut down by inhibiting a selective group of frontal areas simultaneously (bottom row, inhibition to areas 9/46v, 9/46d, F7, and 8B). The shut-down input, of strength I=0.3 and 1s duration, is provided to the nonselective inhibitory population of each of these four areas.

In addition to permanently inactivating areas, we tested the effects of brief (500 ms ∼1s) inactivation in specific areas, and compare the effects in localized vs distributed WM scenarios. For networks relying on localized WM, areas at the top of the hierarchy maintained their selective information largely independent from each other. Consequently, briefly inactivating area 9/46d would not, for example, have an effect on the persistent activity of other areas such as 8B (Fig. 7c, top row). Furthermore, the brief inactivation was enough to remove the information permanently from 9/46d, which remained in the spontaneous state after the inhibition was withdrawn. On the other hand, silencing an area like 9/46d will slightly affect the persistent activity in other areas (such as 8B) in a network strongly relying on distributed WM (Fig. 7c, middle row). However, area 9/46d will be able to recover the encoded selective information once the inhibitory pulse is removed, due to the strong interaction between cortical areas during the delay period. This constitutes a strong prediction for networks which rely on distributed interactions to maintain WM.

The marked resilience of distributed WM attractors to brief inhibitory pulses raises the question of how to shut down the persistent activity once the task has been done. In many traditional WM models, this is achieved by providing a strong inhibitory input to the whole WM circuit, which drives the activity of the selective populations back to their spontaneous state^9,10,33^. It is, however, unrealistic to expect that this approach could also be used for shutting down distributed WM patterns, as it would require a large-scale synchronous inhibitory pulse to all active areas.

We therefore explore in our model whether more spatially selective signals can shut down distributed patterns of activity. In spite of their robustness to sensory distractors as discussed above, we find that distributed WM activity patterns can be shut down with an excitatory input targeting inhibitory populations of areas high in the hierarchy. Fig. 7c (bottom row) shows how a visually evoked distributed WM attractor is deactivated when we deliver excitatory input to the inhibitory populations in the top four areas of the hierarchy (9/46v, 9/46d, F7 and 8B). These prefrontal areas are spatially localized and thought to be highly engaged in WM maintenance, and therefore they are suitable candidates to control the suppression of persistent activity in other cortical areas, such as areas LIP and 24c. Therefore, in spite of engaging cortical areas across all four lobes, distributed WM attractors can be controlled and deactivated by localized inhibition to a small set of frontal areas.

Finally, the distributed nature of WM has also implications for the impact of distractors and similar sensory perturbations on maintenance of selective activity and overall performance of the network. We simulated a delayed response task with distractors (Fig. 8a), in which stimulus A is cued to be maintained in WM and stimulus B is presented as distractor during the delay period (and vice versa). When simulated in the localized WM network, we observed that distractors with the same saliency than the original cue were sufficient to switch the network into the new stimuli, rendering the network easy to distract^10^ (Fig. 8b). For the case of distributed WM, however, the network was highly resilient, and distractors with similar saliency levels as the input cues were filtered out so that working memory storage is preserved (Fig. 8c). Overall, we found that localized WM networks can be distracted with stimuli similar or even weaker than the minimum cue input strength required to encode a WM pattern, while effective distractors need to be about three times as strong in the case of distributed WM networks (Fig. 8d). This difference is due to the robustness of a distributed attractor compared to a local circuit mechanism, but also to the effect of the counterstream inhibitory bias which dampens the propagation of distractor signals (cf. MT responses in Fig. 8b and c). This constitutes a key difference between distributed and local WM models feasible of experimental validation.

**Figure 8:**
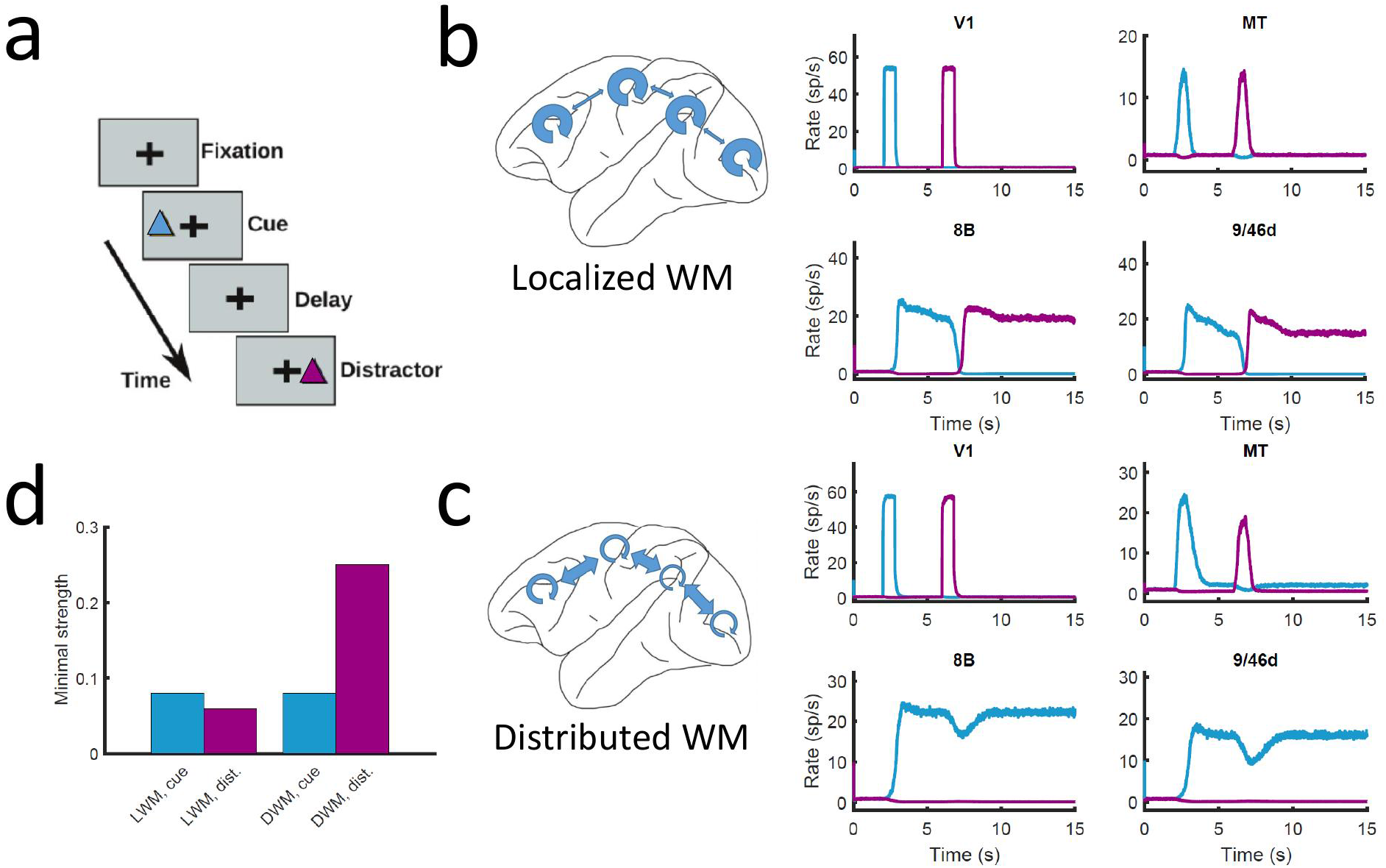
Resistance to distractors in localized vs distributed WM. (a) Scheme of the WM task with a distractor, with the cue (current pulse of strength I_A_=0.3 and duration 500 ms) preceding the distractor (I_B_=0.3, 500 ms) by four seconds. (b) Activity traces of selected areas during the task, for a network displaying localized WM (J_max_=0.468, G=0.21). (c) Same as (b), but for a model displaying distributed WM (J_max_=0.26, G=0.48). (d) Minimal strength required by the cue (blue) to elicit a persistent activity state, and minimal strength required by the distractor (purple) to remove the persistent activity, both for localized WM (left) and distributed WM (right).

## Discussion

The investigation of cognitive functions has been traditionally restricted to operations in local brain circuits –mostly due to the limitations on available precision recording techniques to local brain regions, a problem that recent developments are starting to overcome^1,2^. It is therefore imperative to advance in the study of distributed cognition using computational models as well, to support experimental advances. In this work, we have presented a large-scale circuit mechanism of distributed working memory, realized by virtue of a new concept of robust bifurcation in space. The distributed WM scenario is compatible with recent observations of multiple cortical areas participating in WM tasks^12–14^, even when some of these areas have not been traditionally associated with WM.

One of the main ingredients of the model is the gradient of excitation across the cortical hierarchy, implemented via an increase of excitatory recurrent connections (hinted by the existing anatomical evidence on dendritic spines on pyramidal cells across multiple cortical areas^24^). Moreover, we introduce the concept of counterstream inhibitory bias which was found to stabilize distributed yet spatially confined persistent activity patterns in spite of dense inter-areal connectivity. Macroscopic gradients and hierarchical structures has recently been proposed as a general principle for understanding heterogeneities in the cortex^23^. It has been shown that gradients of circuit properties in line with hierarchical structures contribute to the emergence of a gradient of time scales across cortex, supporting slow dynamics in prefrontal areas^21,23,31^ (see also Extended Data Fig. 9), and also that a hierarchical organization of functional states could serve as basis for WM-guided decisions and executive control^34,35^. It is possible that structural gradients would play a role not only in other cognitive functions in monkeys, but also in other animals including mice^36^ and humans^37,38^.

Theoretically, the present work is the first to show that graded changes of circuit properties along the cortical hierarchy provides a mechanism to explain qualitatively distinct functions of different cortical areas (whether engaged in working memory). This is reminiscent of the phenomenon mathematically called bifurcation, which denotes the emergence of novel behavior as a result of quantitative property change as a control parameter in a nonlinear dynamical system^39^. Our model displays a novel form of bifurcation across the cortex, which cannot be simply explained by a parameter change laid out spatially by virtue of a macroscopic gradient, because areas are densely connected with each other in a complex large-scale network. Bifurcation in space implies a transition near a few association areas which should exhibit signs of dynamical criticality akin to water near a transition between gas and liquid states. This will be explored further in the future.

Interestingly, the model uncovered a host of distinct persistent activity attractor states, each with its own transition location in the cortical tissue. They are defined by their spatial distributed patterns in the large-scale cortical system, independent of the number of selective neural pools per area (Fig. 5a). Many of these persistent activity states are not produced by stimulation of primary sensory areas. These “hidden” attractor states could serve various forms of internal representations such as those that are not triggered by a particular sensory pathway –or those triggered by sensory input but are encoded differently as memories^40^. The identification of these internal representations in further detail are beyond the scope of the present paper, but uncovering their functional role should be within reach of additional experimental and computational work.

### Extending the model of distributed working memory

The reported model of large-scale cortical networks is, to the best of our knowledge, the first of its kind addressing a cardinal cognitive function in a data-constrained way, and it opens the door for elucidating this and similar complex brain processes in future research. Several avenues may be taken to extend the functionality of the present model. First, it is straightforward to have an arbitrary number of selective neural pools per area^9^, which would increase both the selectivity to sensory inputs and the available number of distributed WM attractors. In that case, more complex connections (not necessarily A to A, B to B, etc.) can be investigated, including a distance-dependent “ring structure”^9,10^ or random connections^41^. Second, the model presented here is limited to 30 cortical areas, and can be expanded to include both additional cortical areas and subcortical structures relevant for working memory (such as thalamic nuclei^42^) as their connectivity data become available. An interesting extension in this sense could involve human connectomics data, to reveal the potential influence of complex network structures in the emergence of distributed distractors^38,43,44^. Third, the model can be improved by incorporating more biological details such as cortical layers^26^, contributions of different neuromodulators, and various types of inhibitory neurons.

### Relationship with other mechanisms of working memory

In the description adopted here, we have considered that working memory was maintained via selective persistent activity underlying attractor dynamics, as in most traditional WM approaches^9,33^. However, the proposed principle for distributed WM, and in particular the hypothesis that WM arises due to cooperation between spatially distributed brain areas via long-range recurrent interactions, is also compatible with other mechanisms^45^. For example, recent work suggests that cortex can generate stable WM representations while relying on time-varying, rather than persistent, firing rates^46–48^. There is no reason to think that the encoding of memory items could not use the complex spatiotemporal interactions between brain areas instead of just local interactions. Likewise, silent WM^48–50^, which relies on short-term synaptic reinforcements to sustain information in WM, could also be implemented in a large-scale, distributed scenario. A large-scale implementation of WM also pairs well with recent hypotheses in which memory selectivity is reached via dynamical flexibility instead of content-based attractors, since the wide number and heterogeneity of long-range projections would reduce connection overlap and alleviate the limit capacity of these models^41^. Frameworks of WM in which oscillations play an active role, for example regarding WM-guided executive control^35^, may benefit from using distributed WM approaches, given the usefulness of previous models of large-scale brain networks to explain oscillatory phenomena in the macaque brain^26^.

### Experimental predictions provided by our model

The distributed WM model presented here yields four experimentally testable predictions in monkey (and potentially rodent) experiments, which can be used to validate our theory. First, the model predicts a positive correlation between the number of areas involved in a WM task and their average firing rate of persistent activity (Fig. 5e). Such relationship should not occur according to models of localized WM, since activity levels would be fairly independent across areas. This prediction could be tested with neuroimaging experiments. A complementary version of this prediction is that, if areas displaying persistent activity are silenced (e.g. optogenetically), the activity of the other persistent activity areas will decrease (Fig. 6b).

Second, our model predicts that areas involved in distributed WM patterns can be briefly silenced without losing the encoded information, which will be recovered as soon as the inhibition is gone (Fig. 7, middle row), something that localized WM do not predict (Fig.7 top row). This also means that silent activity periods associated with silent WM^48–50^ could also be due to distributed WM effects. Optogenetic inactivation could be used to test this result.

Third, distributed WM is significantly more robust to distractors than localized WM (Fig. 8), due to their intrinsic resilience and the inhibitory feedback condition. Behavioral experiments in macaques should be able to test this robustness to distractors.

Fourth, electrophysiological recordings in macaques could test whether FEF areas require support from frontal areas (in the form of strong excitation) to maintain WM-related activity (Extended Data Fig. 7). Although this prediction focuses on a particular set of areas, it should shed light into unclear aspects of FEF dynamics.

In a more general sense, our model predicts a reversed gradient of inhibition and strong large-scale interactions to sustain distributed WM patterns, which may be observed using different experimental approaches. It will also be interesting to see whether the same model is able to account for decision-making processes as well as working memory^20,27^.

Conceptually, this work revealed a novel form of bifurcation in cortical space as a mechanism to generate differential functions across different cortical areas, a concept that is likely to be generalizable for understanding how distinct cortical areas endowed with a canonical circuit organization are at the same time suited for differential functions^23^.

## Online Methods

### Anatomical data

The anatomical connectivity data used has been gathered in an ongoing track tracing study in macaque and has been described in detail elsewhere^17–19,26^. Briefly, retrograde tracer injected into a given target area labels neurons in a number of source areas projecting to the target area. By counting the number of labeled neurons on a given source area, Markov et al. defined the fraction of labeled neurons (FLN) from that source to the target area. FLN can serve as a proxy for the “connection strength” between two cortical areas, which yields the connectivity pattern of the cortical network (Extended Data Fig. 1a, b). In addition, Markov et al. also measured the number of labeled neurons located on the supragranular layer of a given source area. Dividing this number over the total number of labeled neurons on that source area, we can define the supragranular layered neurons (SLN) from that source area to the target area (Extended Data Fig. 1c, d).

SLN values may be used to build a well-defined anatomical hierarchy^19,25^. Source areas located lower (higher) than the target area in the anatomical hierarchy, as defined in^25^, display a progressively higher (lower) proportion of labeled neurons in the supragranular layer. As a consequence, the lower (higher) the source area relative to the target area, the higher (lower) the SLN values of the source-to-target projection. By performing a logistic regression on the SLN data to accommodate each area in its optimal position in the anatomical hierarchy^21^, we assign a hierarchical value h_i_ to each area ‘i’.

Iterating these measurements across other anatomical areas yields and anatomical connectivity matrix with weighted directed connections and an embedded structural hierarchy. The 30 cortical used to build our data-constrained large-scale brain network are, in hierarchical order: V1, V2, V4, DP, MT, 8m, 5, 8l, 2, TEO, F1, STPc, 7A, 46d, 10, 9/46v, 9/46d, F5, TEpd, PBr, 7m, LIP, F2, 7B, ProM, STPi, F7, 8B, STPr and 24c. Finally, data on wiring connectivity distances between cortical areas is available for this dataset as well, allowing to consider communication time lags when necessary (we found however that introducing time lags this way does not have a noticeable impact on the dynamics of our model). The connectivity data used here is available to other researchers from core-nets.org.

The corresponding 30×30 matrices of FLN and SLN are shown in Extended Data Fig. 1b, d. Areas in these matrices are arranged following the anatomical hierarchy, which is computed using the SLN values and a generalized linear model^21,26^. Surgical and histology procedures were in accordance with European requirements 86/609/EEC and approved by the ethics committee of the region Rhone-Alpes.

In addition to the data on FLN and SLN across 30 cortical areas, we used additional data to constrain the area-to-area differences in the large-scale brain network. In particular, we have collected data on the total spine count of layer 2/3 pyramidal neuron basal dendrites across different cortical areas, as the spine count constitutes a proxy for the density of synaptic connections within a given cortical area^24^. A full list of all area-specific values of spine densities considered and their sources is given in Extended Data Table 1.We use an age correction factor meant to correct for the decrease of spine counts with age for data obtained from old monkeys. A plausible estimate would be a ∼30% decrease for a 10y difference^51,52^. See Extended Data Fig. 2 for the effect of this correction on the overall gradient established by the spine count data, and the correlation of such gradient with the SLN hierarchy.

### Experimental evidence of WM-related activity across cortical areas

To compare the results of our model with existing evidence, we generated brain maps highlighting areas for which experimental evidence of WM-related activity during the delay period has been found. Following the data collected by recent review studies^12,13^, we distinguish between three categories. First, areas with strong WM evidence (for which at least two studies show support of WM-related activity, or if only studies supporting WM activity are known) are shown in dark blue in the maps of Figs. 2 and 4. Second, areas with moderate evidence (for which substantial positive and negative evidence exist) are shown in light blue. Finally, areas for which strong negative evidence exists (more than two studies with negative evidence, or absence of any positive studies) are left as grey in the map. Alternative criteria have only small effects on the resulting maps and the general results are consistent to variations.

### Computational model: Local neural circuit

We describe the neural dynamics of the local microcircuit representing a cortical area with the Wong-Wang model^27^. In its three-variable version, this model describes the temporal evolution of the firing rate of two input-selective excitatory populations as well as the evolution of the firing rate of an inhibitory population. All populations are connected to each other (see Fig. 1a). The model is described by the following equations:

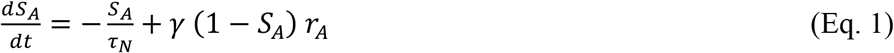

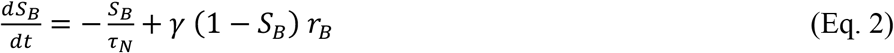

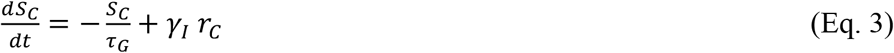

Here, S_A_ and S_B_ are the NMDA conductances of selective excitatory populations A and B respectively, and S_C_ is the GABAergic conductance of the inhibitory population. Values for the constants are τ_N_=60 ms, τ_G_=5 ms, γ=1.282 and γ_I_=2. The variables r_A_, r_B_ and r_C_ are the mean firing rates of the two excitatory and one inhibitory populations, respectively. They are obtained by solving, at each time step, the transcendental equation *r*_*i*_ = *ϕ*_*i*_(*I*_*i*_) (where *ϕ* is the transfer function of the population, detailed below), with I_i_ being the input to population ‘i’, given by

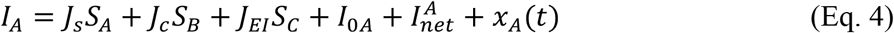

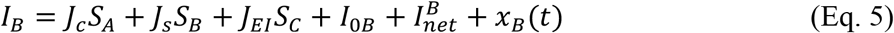

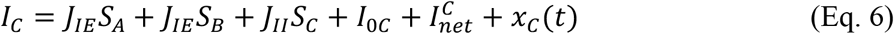

In these expressions, J_s_, J_c_ are the self- and cross-coupling between excitatory populations, respectively, J_EI_ is the coupling from the inhibitory populations to any of the excitatory ones, J_IE_ is the coupling from any of the excitatory populations to the inhibitory one, and J_II_ is the self-coupling strength of the inhibitory population. The parameters I_0i_ with i=A, B, C are background inputs to each population. Parameters are J_s_=0.3213 nA, J_c_=0.0107 nA, J_IE_=0.15 nA, J_EI_=-0.31 nA, J_II_=-0.12 nA, I_0A_=I_0B_=0.3294 nA and I_0C_=0.26 nA. Later we will modify some of these parameters in an area-specific manner (in particular J_s_ and J_IE_) to introduce a gradient of properties across the cortical hierarchy. The term I^i^_net_ denotes the long-range input coming from other areas in the network, which we will keep as zero for now but will be detailed later. Sensory stimulation can be introduced here as extra pulse currents of strength I_pulse_ =0.3 and duration T_pulse_ =0.5 sec (unless specified otherwise).

The last term x_i_(t) with i=A, B, C is an Ornstein-Uhlenbeck process, which introduces some level of stochasticity in the system. It is given by

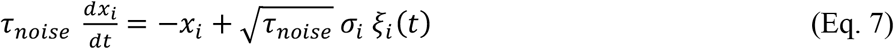

Here, ξ_i_(t) is a Gaussian white noise, the time constant is τ_noise_=2 ms and the noise strength is σ_A,B_=0.005 nA for excitatory populations and σ_C_=0 for the inhibitory one.

The transfer function ϕ_i_(t) which transform the input into firing rates takes the following form for the excitatory populations^53^:

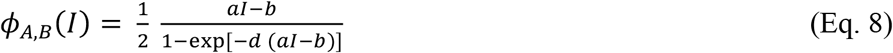

The values for the parameters are *a*=135 Hz/nA, *b*=54 Hz and *d*=0.308 s. For the inhibitory population a similar function can be used, but for convenience we choose a threshold-linear function:

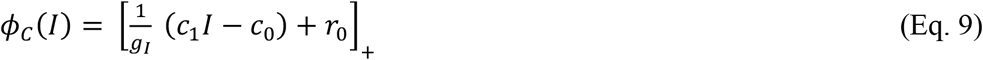

The notation [*x*]_+_ denotes rectification. The values for the parameters are g_I_=4, c_1_=615 Hz/nA, c_0_=177 Hz and r_0_=5.5 Hz. Finally, it is sometimes useful for simulations (although not a requirement) to replace the transcendental equation *r*_*i*_ = *ϕ*_*i*_(*I*_*i*_) by its analogous differential equation, of the form

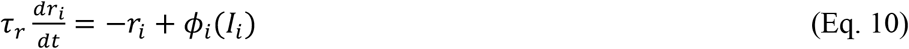

The time constant can take a typical value of τ_r_=2 ms.

### Computational model: Gradient of synaptic strengths

Before considering the large-scale network and the inter-areal connections, we look into the area-to-area heterogeneity to be included in the model.

Our large-scale cortical system consists of N=30 local cortical areas, for which inter-areal connectivity data is available. Each cortical area is described as a Wong-Wang model of three populations like the ones described in the previous section. Instead of assuming areas to be identical to each other, here we will consider some of the natural area-to-area heterogeneity that has been found in anatomical studies. For example, work from Elston^24^ has identified a gradient of dendritic spine density, from low spine numbers found in early sensory areas to large spine counts found in higher cognitive areas. This may reflect an increase of local recurrent strength as we move from sensory to association areas. In addition, cortical areas are distributed along an anatomical hierarchy^19,25^. The position of a given area ‘i’ within this hierarchy, namely h_i_, can be computed via a logistic regression of the SLN (fraction of supragranular layer neurons) projecting to and from that area^21^.

In the following, we will assign the incoming synaptic strength (both local and long-range) of a given area as a linear function of the dendritic spine count values observed in anatomical studies, with age-related corrections when necessary. Alternatively, when spine count data is not available for a given area, we will use its position in the anatomical hierarchy, which displays a high correlation with the spine count data, as a proxy for the latter. After this process, the large-scale network will display a gradient of local and long-range recurrent strength, with sensory/association areas showing weak/strong local connectivity, respectively. We denote the local and long-range strength value of a given area *i* in this gradient as h_i_, and this value normalized between zero (bottom of the gradient, area V1) and one. In summary:

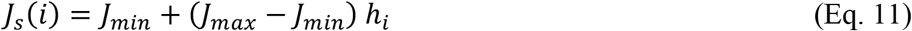

We assume therefore a gradient of values of J_s_, with its value going from J_min_ to J_max_. Having large values of J_s_ for association areas affects the spontaneous activity of these areas, even without considering inter-areal coupling. A good way to keep the spontaneous firing rate of these areas within physiologically realistic limits is to impose that the spontaneous activity fixed point is the same for all areas^16^. To introduce this into the model, we take into account that the solutions in the spontaneous state are symmetrical: S_A_=S_B_=S (we assume zero noise for simplicity). The current entering any of the excitatory populations is then (assuming I_0A_=I_0B_=I_0_):

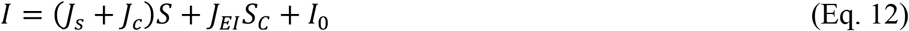

Assuming a fast dynamics for r_C_ and S_C_ (mediated by GABA) as compared to S_A_ and S_B_ (mediated by NMDA) we can obtain the approximate expression for S_C_:

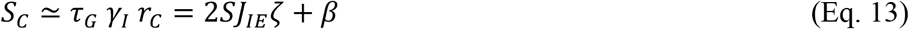

with

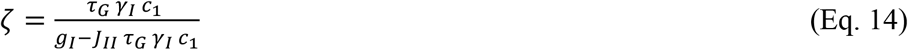

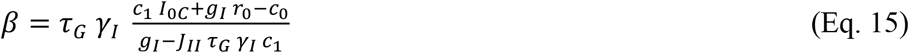

The equation for the excitatory current has then the form

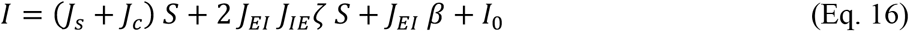

To maintain the excitatory input (and therefore the spontaneous activity level S) constant while varying J_s_ across areas, we just have to keep the quantity *J*_*s*_ + *J*_*c*_ + 2 *J*_*EI*_ *J*_*IE*_ *ζ* ≡ *J*_0_ constant (for the original parameters of the isolated area described above, we obtain J_0_=0.2112 nA). A good choice, but not the only one, is to assume that the excitatory synapses to inhibitory neurons, J_IE_, also scales with the ranks and with J_s_ accordingly:

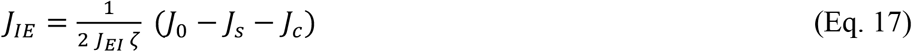

This linear relationship ensures that the spontaneous solution is the same for all areas in the network. Note that deviations from this linear relationship would simply lead to different areas having slightly different spontaneous activity levels, but it does not substantially affect our main results.

Since J_IE_ needs to be non-negative, the linear relationship above imposes a minimum value of J_min_=0.205 nA for J_s_. The particular maximum value of J_s_, namely J_max_, will determine the type of WM model we assume. Since the bifurcation point of an isolated area is at J_s_=0.4655 nA for this set of parameter values, setting J_max_ below that value implies that all areas in the network are monostable in isolation. In this situation, any persistent activity displayed by the model will be a consequence of a global, cooperative effect due to inter-areal interactions. On the other hand, having J_max_ above the bifurcation point means that some areas will be multistable when isolated, e.g. they will be intrinsically multistable and compatible with classical WM theories.

Unless specified otherwise, we assume a range of J_min_=0.21 nA and J_max_=0.42 nA (i.e. below the critical value), so that the model displays distributed WM without having any intrinsically bistable areas.

### Computational model: Inter-areal projections

We now consider the inter-areal projections connecting isolated areas to form the large-scale cortical network. Assuming that inter-areal projections stem only from excitatory neurons (as inhibitory projections tend to be local in real circuits) and that such projections are selective for excitatory neurons, the network or long-range input term arriving at each of the populations of a given area *x* from all other cortical areas is given by

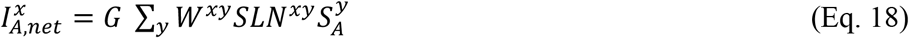

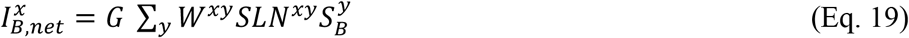

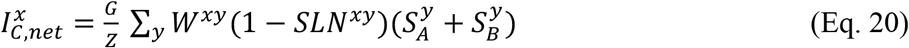

Here, G is the global coupling strength, Z is a balancing factor, and W is the connectivity matrix (more details given below). In these equations, a superindex denotes the cortical area and a subindex the particular population within each area. The sum in all equations runs over all cortical areas of the network (N=30). Excitatory populations A and B receive long-range inputs from equally selective units from other areas, while inhibitory populations receive inputs from both excitatory populations. Therefore, neurons in population A of a given area may be influenced by A-selective neurons of other areas directly, and by B-selective neurons of other areas indirectly, via local interneurons.

G is the global coupling strength, which controls the overall long-range projection strength in the network (G=0.48 unless specified otherwise). Z is a factor that takes into account the relative balance between long-range excitatory and inhibitory projections. Setting Z=1 means that both excitatory and inhibitory long-range projections are equally strong, but this does not guarantee that their effect is balanced in the target area, due to the effect of local connections. Following previous work^16^, we choose to impose a balance condition that guarantees that, if populations A and B have the same activity level, their net effect on other areas will be zero –therefore highlighting the selectivity aspect of the circuits. Again, deviations from this balance condition do not strongly affect our results besides the appearance of small differences between populations A and B. Considering that the transfer function of inhibitory populations is linear and their approximately linear rate-conductance relationship, it can be shown that

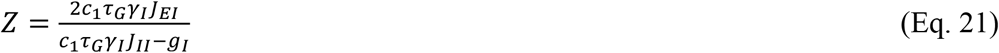

Aside from global scaling factors, the effect of long-range projections from population *y* to population *x* is influenced by two factors. The first one, *W*^*xy*^, is the anatomical projection strength as revealed by tract-tracing data^17^. We use the fraction of labelled neurons (FLN) from population *y* to *x* to constrain our projections values to anatomical data. We rescale these strengths to translate the broad range of FLN values (over five orders of magnitude) to a range more suitable for our firing rate models. We use a rescaling that maintains the proportions between projection strengths, and therefore the anatomical information, that reads

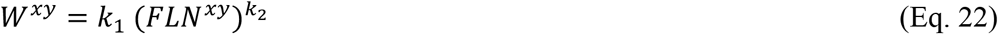

Here, the values of the rescaling are *k*_*1*_ =1.2 and *k*_*2*_ =0.3. The same qualitative behavior can be obtained from the model if other parameter values, or other rescaling functions, are used as long as the network is set into a standard working regime (i.e. signals propagate across areas, global synchronization is avoided, etc.) FLN values are also normalized so that ∑_*y*_ *FLN*^*xy*^ = 1. While in-degree heterogeneity might impact network dynamics^54,55^, this was done to have a better control of the heterogeneity levels of each area, and to minimize confounding factors such as the uncertainty on volume injections of tract tracing experiments and the influence of potential homeostatic mechanisms. In addition, and as done for the local connections, we introduce a gradient of long-range projection strengths using the spine count data: *W*^*xy*^ → (*J*_*s*_(*x*)/ *J*_max_) *W*^*xy*^, so that long-range projections display the same gradient as the local connectivity presented above.

The second factor that needs to be taken into account is the directionality of signal propagation across the hierarchy. Feedforward (FF) projections that are preferentially excitatory constitute a reasonable assumption which facilitate signal transmission from sensory to higher areas. On the other hand, having feedback (FB) projections with a preferential inhibitory nature contributes to the emergence of realistic distributed WM patterns (Fig. 4) (see also previous work^19,30^). This feature can be introduced, in a gradual manner, by linking the different inter-areal projections with the SLN data, which provides a proxy for the FF/FB nature of a projection (SLN=1 means purely FF, and SLN=0 means purely FB). In the model, we assume a linear dependence with SNL for projections to excitatory populations and with (1-SLN) for projections to inhibitory populations, as shown above.

Following recent evidence of frontal networks having primarily strong excitatory loops^56^, it is convenient to ensure that the SLN-driven modulation of FB projections between frontal areas is not too large, so that interactions between these areas are never strongly inhibitory. In practice, such constraint is only necessary for projections from frontal areas to 8l and 8m (which are part of the frontal eye fields) and has little effect on the behavior of our model otherwise. The introduction of this limitation has two minor consequences: (i) it allows area 8l and 8m to exhibit a higher level of persistent activity during distributed WM –as their hierarchical position and recurrent strength are not strong enough to sustain activity otherwise, as previously suggested in anatomical studies^17,19^, and (ii) it slightly shifts the bifurcation point in cortical space (see Extended Data Fig. 7). Unless specified otherwise (and in Fig. 4, where the limitation is not considered for a cleaner study of the effects of inhibitory feedback), we consider that the SLN-driven modulation of FB projections to 8l and 8m is never larger than 0.4.

### Simplified computational model: two areas

To provide a deeper intuition of the nature of distributed WM and the model ingredients which are fundamental for the phenomenon, we describe here a simplified version of our model which is suitable for theoretical analysis. We will first introduce a version of the model with two interconnected excitatory nodes, each of them following a rate dynamics:

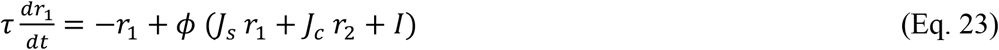

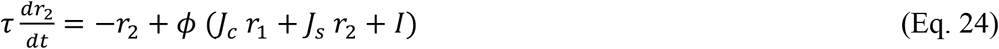

Here, *r*_1_, *r*_2_ are the firing rates of node 1 and 2, with display self-connections of strength *J*_*s*_ and cross-connections of strength *J*_*c*_. Both areas have a relaxation time constant *τ* and receive a background input current *I*. The transfer function is a threshold-linear function, i.e. *ϕ*(*I*) = *I* − *θ* if *I* > *θ* and zero otherwise.

The equation for the fixed point of the dynamics can be easily written as a function of the average firing rate of both units, namely R, which leads to the equation

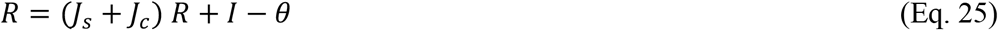

The above is only true in the linear regime, while for the subthreshold regime we get the trivial solution R=0. Solving Eq. 25 leads to the following expression

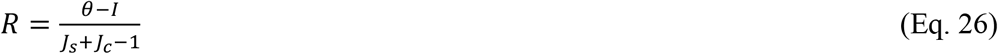

Eq. 26 permits a positive solution for R if both numerator and denominator are positive (or both negative, but the solution in this case is less interesting biologically). Interestingly, if the cross-connection strength is large enough, we can have a fixed point solution with nonzero R even if *J*_*s*_ < 1, which would correspond to the case in which nodes would be monostable if isolated from each other. Distributed persistent activity can emerge, therefore, even in simple cases of two coupled excitatory nodes.

### Simplified computational model: N areas

We present now the case in which we have a large number N of connected excitatory nodes. We consider a fully connected network for simplicity and N=30 nodes for the simulations. The activity of the nodes is given by

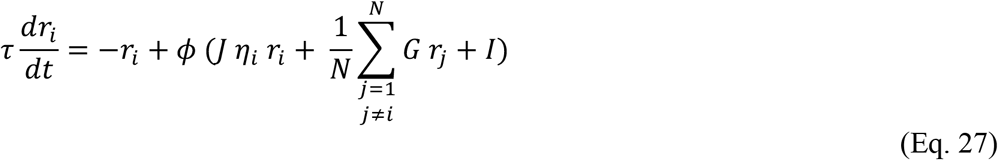

Here, the term *η*_*i*_ is a monotonically increasing linear function that is used to introduce a gradient of connectivity strength across the network, with minimum value *η*_1_ = 0.55 and maximum value *η*_*N*_ = 0.85. Other parameters values chosen for simulations are: *τ* = 20 *ms, I* = 4.81, *J* = 0.91. For the transfer function, we choose a sigmoidal function (to demonstrate robustness of the phenomenon observed before for the threshold-linear):

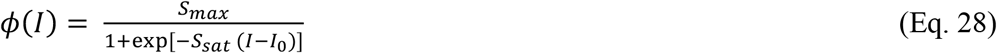

Parameters chosen for simulations are *S*_*max*_ = 60, *S*_*sat*_ = 0.1 *I*_0_ = 30. For these parameters, an isolated node is bistable only if *η* ≅ 0.88, which is above our chosen *η*_*N*_ value and implies that all nodes are monostable in isolation.

This model admits an approximate mean-field solution if we define the network-average firing rate as 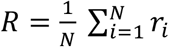. Averaging Eq. 27 over units and using standard mean-field approximations like ⟨*f*(*x*)⟩ ≈ *f*(⟨*x*⟩), we arrive at

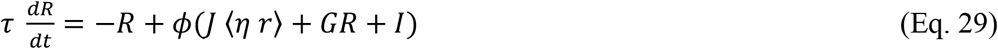

To estimate the average of the product *η r* over units, we can follow to approaches. First, we can assume independence between these two variables and accept that ⟨*η r*⟩ ≈ ⟨*η*⟩ ⟨*r*⟩ = *η*_0_ *R*, with *η*_0_ being the mean value of *η*. We will refer to this as the first-order mean field solution in the text, and is given by the following equations:

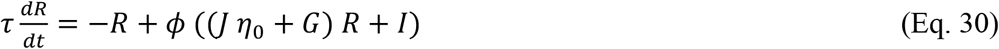

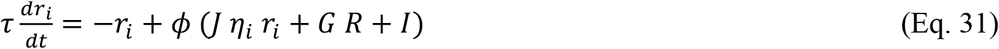

It is important to notice that Eq. 30 is the real self-consistent mean-field solution, which can be solved numerically to find the fixed point solutions of our system. Eq. 31, on the other hand, is useful since it allows to obtain the fixed point solutions for any node ‘i’ that is connected to the network, and allows to explore the effect of the local heterogeneity *η*.

As an alternative to this solution, we can consider additional term in the dependence between the heterogeneity and firing rate of the nodes. For example, we can assume that the firing rate of units will grow proportionally to both *η* and G, which is a plausible approximation that takes into account both the heterogeneity and the global interactions. We can write this as

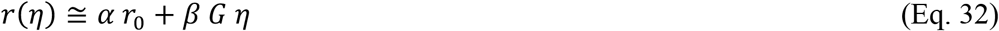

This leads to the following expression:

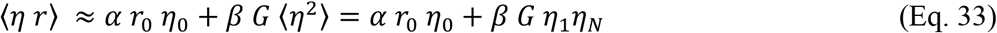

The mean-field solution obtained, which we denote here as second-order solution, is given by

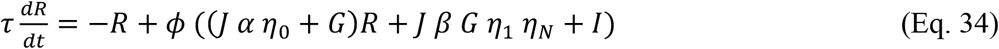

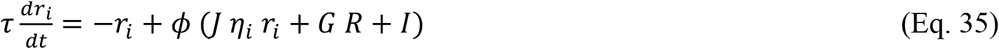

A choice of *α* = 0.94 and *β* = 10 (which fulfills the recommendations for a perturbative approach, since α ≈ 1 and *βη*_1_*η*_*N*_ ≪ 1) provides good results as Fig. 3 shows. However, both the first and second order solutions predict the emergence of distributed WM, and the choice of one or the other solution has only minor implications.

### Data analysis: Overview

We developed a numerical method to estimate the number of stable distributed WM attractors for a particular set of parameters values of our model. This method, which follows simplified density-based clustering principles, is used to obtain the results shown in Figs. 5 and 6. To allow for a cleaner estimation, we do not consider noise in the neural dynamics during these simulations.

Our large-scale cortical network has 30 areas, with each of them having two selective excitatory populations A and B. Simply assuming that each of the areas can reach one of three possible states (persistent activity in A, persistent activity in B, or spontaneous activity) means that our model can potentially display up to 3 to the power of 30 attractor combinations. This number can be even larger if we refine the firing rate reached by each area rather than simply its persistent/non-persistent activity status. Since it is not possible to fully explore this extremely large number of possible attractors, we devised a strategy based on the exploration of a sample of the input space of the model. The core idea is to stimulate the model with a certain input pattern (targeting randomized areas) and registering the fixed point that the dynamics of the model converges to. By repeating this process with a large number of input combinations and later counting the number of different attractors from the obtained pool of fixed points, we can obtain an estimate of the number of attractors for a particular set of parameter values.

### Data analysis: Stimulation protocol

A given input pattern is defined as a current pulse of fixed strength (I_pulse_ =0.2) and duration (T_pulse_ =1 sec) which reaches a certain number P of cortical areas. Only one population (A or B, randomized) in each area receives the input, and the P cortical areas receiving the input are randomly selected across the top 16 areas of the spine count gradient. This decreases the amount of potential input combinations we have to deal with by acknowledging that areas with stronger recurrent connections (such as 9/46d) are more likely to be involved in distributed WM patterns than those with weaker connections (such as MT). P can take any value between one and P_max_ =16, and we run a certain number of trials (see below) for each of them. Different values of I_pulse_ and T_pulse_, as well as setting the randomly selected areas at a high rate initial condition instead of providing an external input, have been also explored and lead to qualitatively similar results. Similar approaches regarding the use of random perturbations in high dimensional systems have been successfully used in the past^57^.

It is also important to consider that not all values of P have the same number of input combinations. For example, P=1 allows for 16*2=32 different input combinations (if we discriminate between populations A and B), while P=2 allows for 16*(16-1)*2=480 input combinations, and so on. For a given value of P, the number of possible input combinations N_c_ is given by

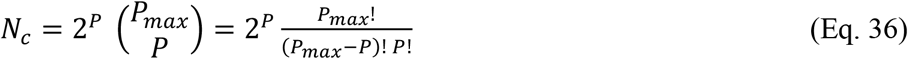

By summing all values of N_c_ for P=1, …P_max_, we obtain around 43 million input combinations, which are still too many trials to simulate for a single model configuration. To simplify this further, we consider a scaling factor F_c_ on top of N_c_ to bring down these numbers to reasonable levels for simulations. We use F_c_ =0.0002 (or 0.02% of all possible combinations) for our calculations, which brings down the total number of simulated input combinations to around 9000. Other options, such as decreasing P_max_ and using a larger scaling factor (P_max_ =12, F_c_ =0.01 or 1% or all possible combinations) give also good results. Since the rescaling can have a strong impact for small P (yielding a number of trials smaller than one), we ensure at least one trial for these cases.

To guarantee the stability of the fixed points obtained during these simulations, we simulate the system during a time windows of 30 seconds (which is much larger than any other time scale in the system), and check that the firing rates have not fluctuated during the last 10 seconds before we register the final state of the system as a fixed point.

### Data analysis: Estimating the number of attractors

The final step is to count how many different attractors have been reached by the system, by analyzing the pool of fixed points obtained from simulations. A simple way to do this is to consider that, for any fixed point, the state of each area can be classified as persistent activity in population A (i.e. mean firing rate above a certain threshold of 10 spikes/s), persistent activity in population B, or spontaneous activity (both A and B are below 10 spikes/s). This turns each fixed point into a vector of 30 discrete states, and the number of unique vectors among the pool of fixed points can be quickly obtained using standard numerical routines in Matlab (such as the ‘unique’ function).

A more refined way to count the number of attractors, which we use in this work, is to define the Euclidean distance to discriminate between an attractor candidate and any previously identified attractors. Once the first attractor (i.e. the first fixed point analyzed) is identified, we test whether the next fixed point is the same than the first one by computing the Euclidean distance E_d_ between them:

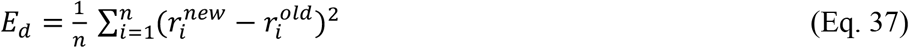

where n=30 is the total number of areas in the network (only one of the populations, A or B, needs to be considered here). If E_d_ is larger than a certain threshold distance ε, we consider it a new attractor. We choose ε=0.01, which grossly means that two fixed points are considered as different attractors if, for example, the activity of one of their cortical areas differs by 0.5 spikes/s and the activity on all other areas is the same for both. The particular value of ε does not have a strong impact on the results (aside from the fact that smaller values of ε gives us more resolution to find attractors). When several attractors are identified, each new candidate is compared to all of them using the same method.

Both the first and the second method to count attractors deliver qualitatively similar results (in terms of the dependence of the number of attractors with model parameters), although as expected the second method yields larger numbers due to its higher discriminability.

## Acknowledgments

We thank Rishidev Chaudhuri, John Murray and Jorge Jaramillo for their support during the development of this work, and Henry Kennedy for providing the connectivity dataset.

## Funding

This work was supported by the NIH grant R01MH062349 (XJW), ONR grant N00014-17-1-2041 (XJW), the Simons Foundation Collaboration on the Global Brain grant (XJW), the Human Brain Project (Flagship, project id. 785907, JFM) and the University of Amsterdam (JFM).

## Author contributions

JFM and X-JW designed the study, JFM performed the research, JFM and X-JW discussed the results and wrote the manuscript.

## Competing interests

Authors declare not competing interests.

## Data and materials availability

All information needed to reproduce the results of this manuscript are in the main text and supplementary material, and the code used will be made available upon publication of this work.

## Supplementary Information

**Extended Data Fig. 1:**
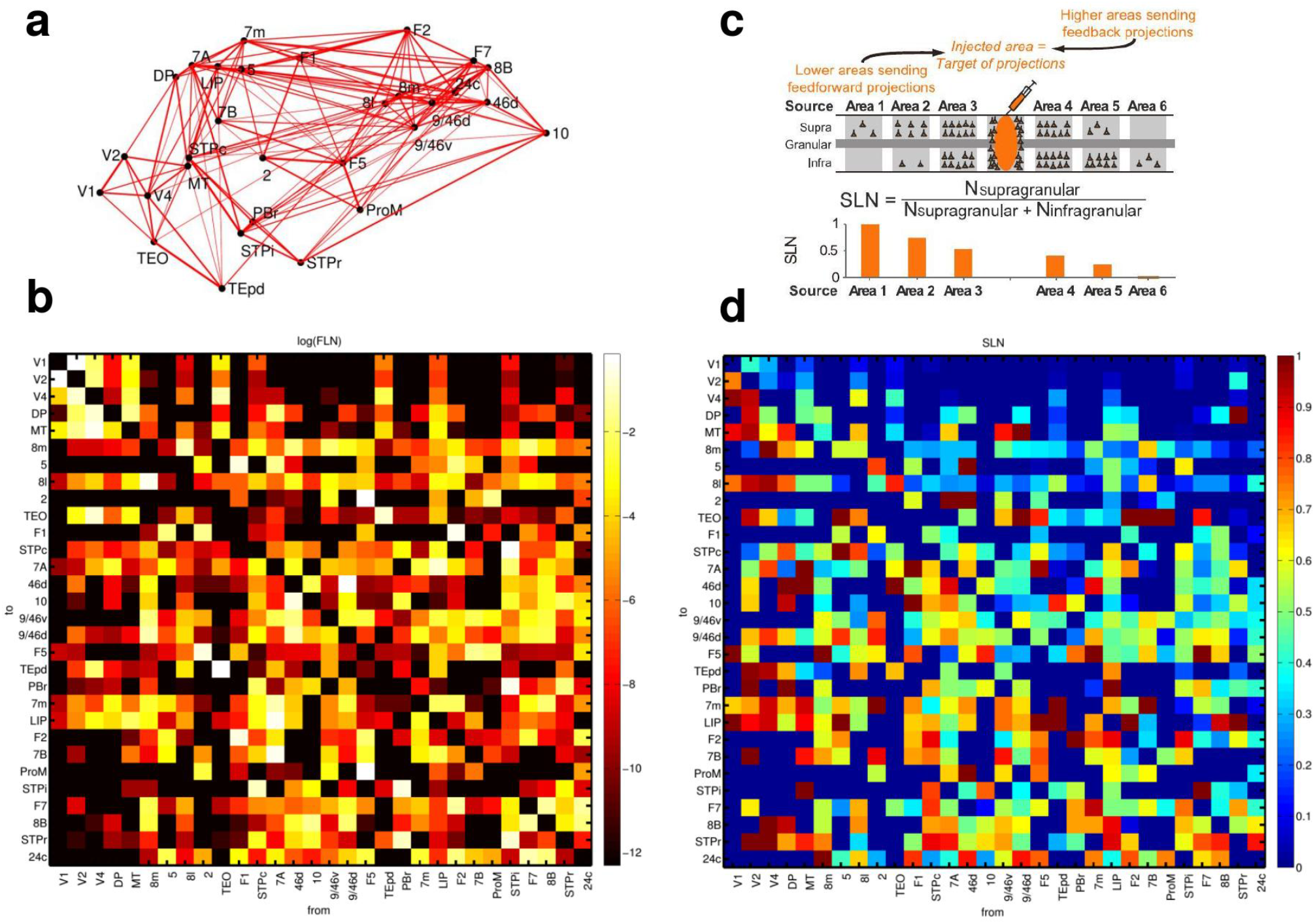
Anatomical connectivity data of the macaque cortex. Data from anatomical studies^**17–19**^. (a) Connectivity of the 30 areas (positioned in 3D space following injection coordinates of experiments). Width of the lines denote two-way averaged FLN values (i.e. average strength of the projection). (b) Map of FLN values for all connections considered. (c) The proportion of supragranular vs infragranular neurons projecting to the injection site allowed to define an anatomical hierarchy. (d) Map of SLN values for all connections considered.

**Extended Data Fig. 2:**
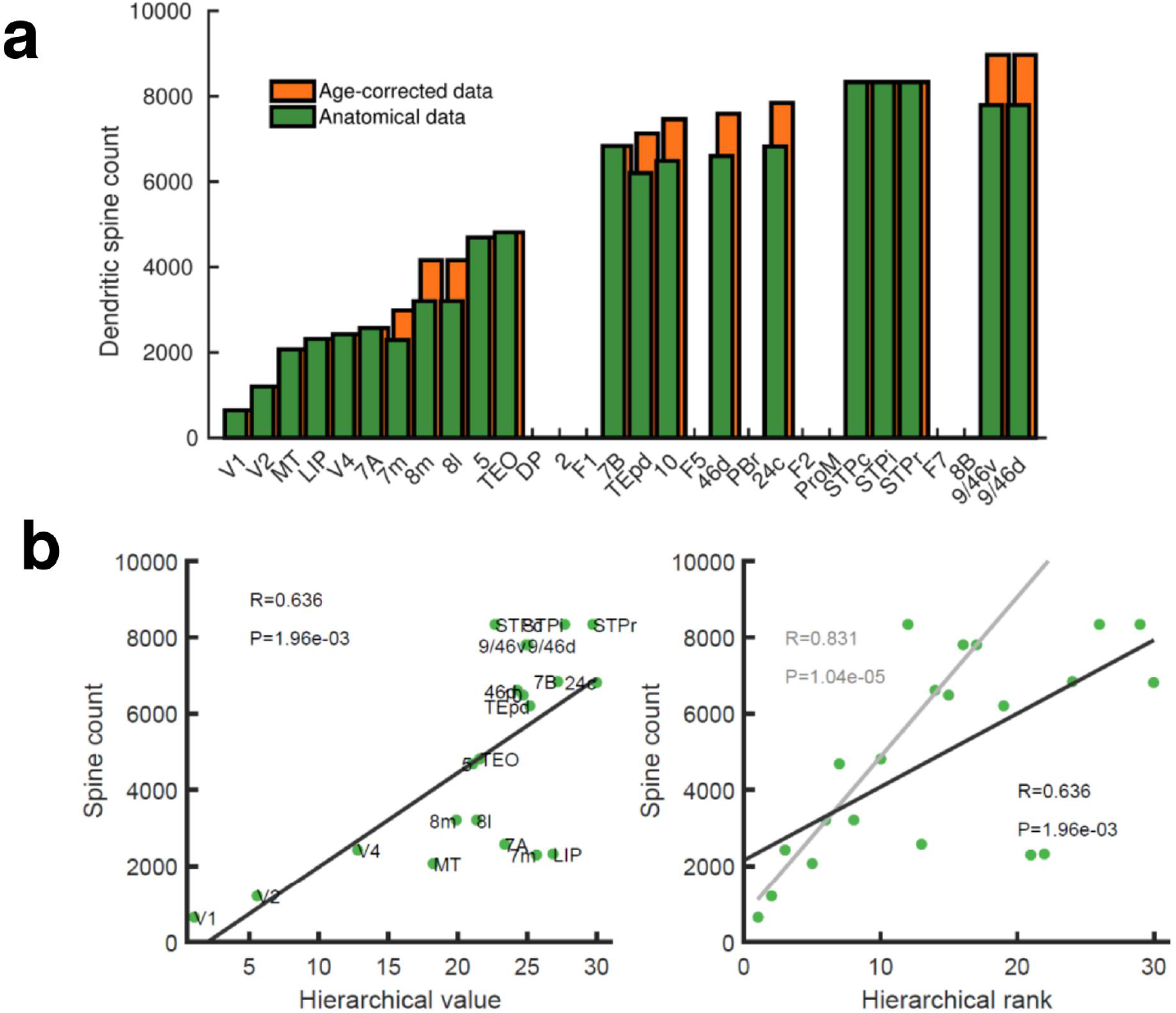
Spine count data used to constrain connectivity strength. Data from anatomical studies^**24**^, see also Extended Data Table 1. (a) Spine count of the basal dendrites of layer 2/3 neurons across cortical areas of young (2∼3 years old) macaques. When data from older macaques had to be considered, an age correction was introduced (orange bars). (b) Correlation between the spine count data and the hierarchical value h_i_ (left) or the hierarchical position/rank (right). A robust fit obtained by random sampling consensus (grey, 100 randomized trials with a threshold for number of outliers <10% of the total points) suggest areas 7m and LIP as potential outliers. Pearson correlation and corresponding P-value are shown in each panel. In the model, the connectivity strength of areas for which spine count data was not available was estimated using their hierarchical value.

**Extended Data Fig. 3:**
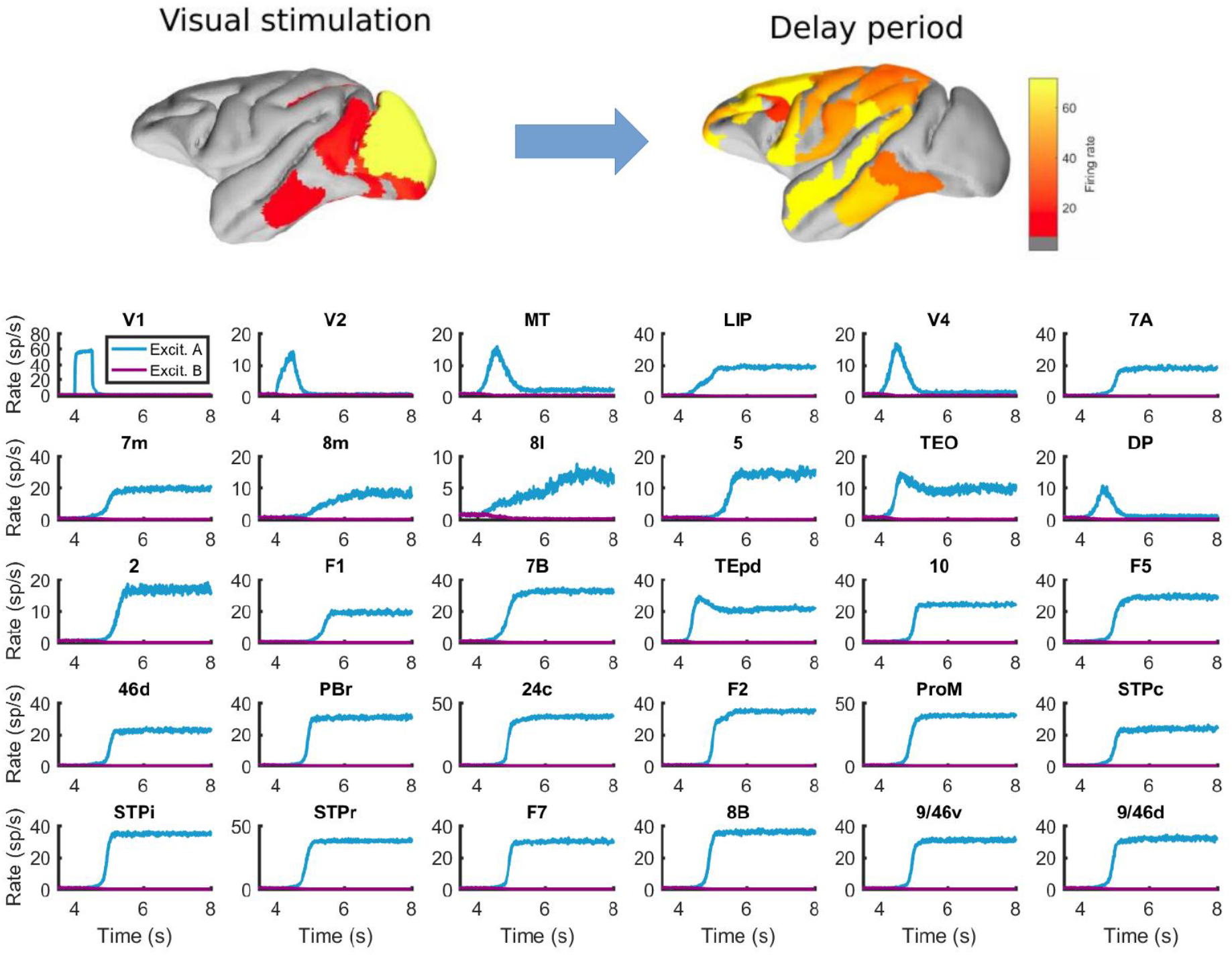
Behavior of all areas in the network during the visual WM task. Top: spatial maps of the simulated macaque brain during stimulation and delay period, with activity color coded. Bottom: evolution of the firing rate of all areas in the network. Stimulation occurs in V1 at t=4 seconds and has a duration of 500 ms. Parameters as in Fig. 2.

**Extended Data Fig. 4:**
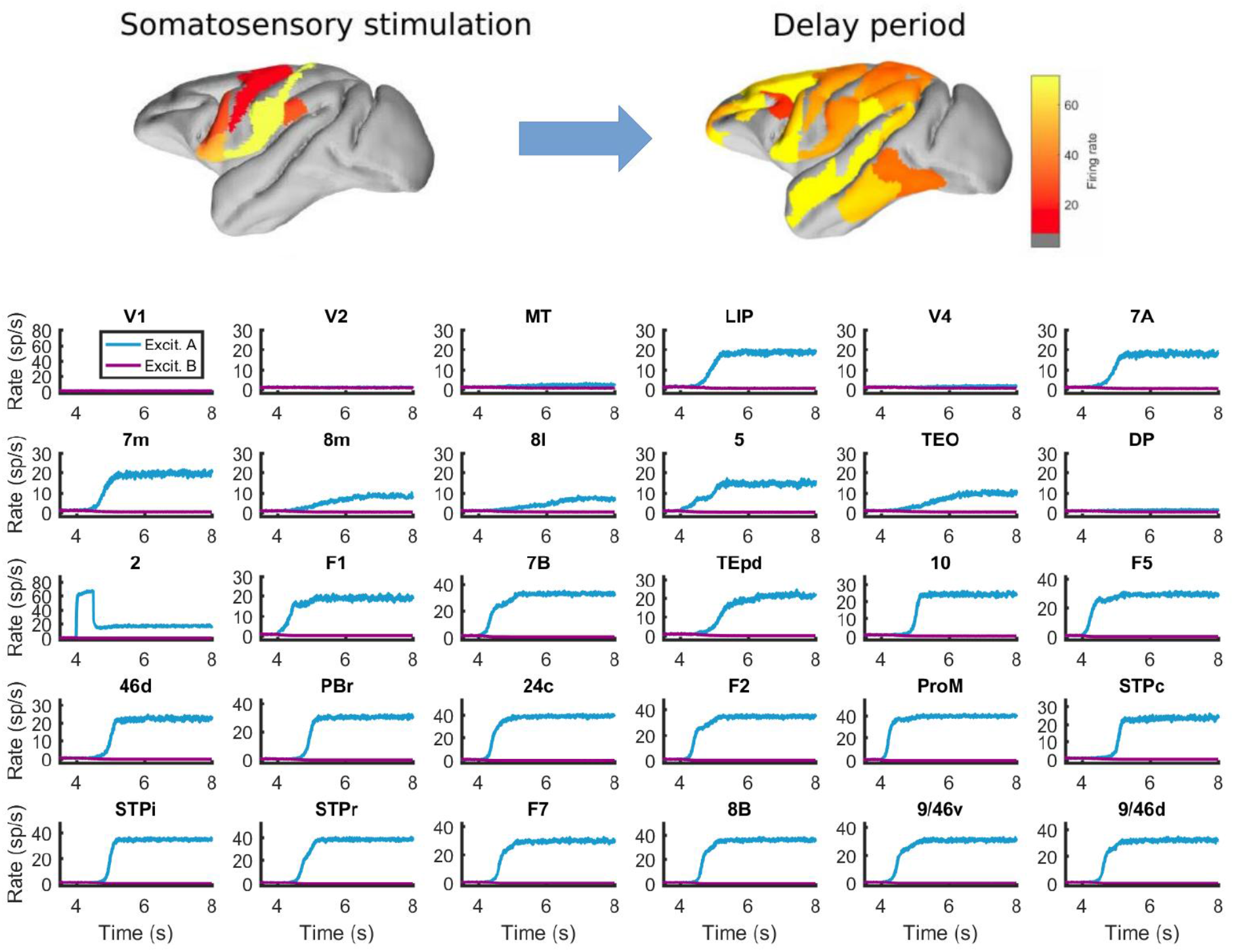
Behavior of all areas in the network during the somatosensory WM task. Top: spatial maps of the simulated macaque brain during stimulation and delay period, with activity color coded. Bottom: evolution of the firing rate of all areas in the network. Stimulation occurs in area 2 (primary somatosensory area) at t=4 seconds, and has a duration of 500ms. Other parameters as in Fig. 2.

**Extended Data Fig. 5:**
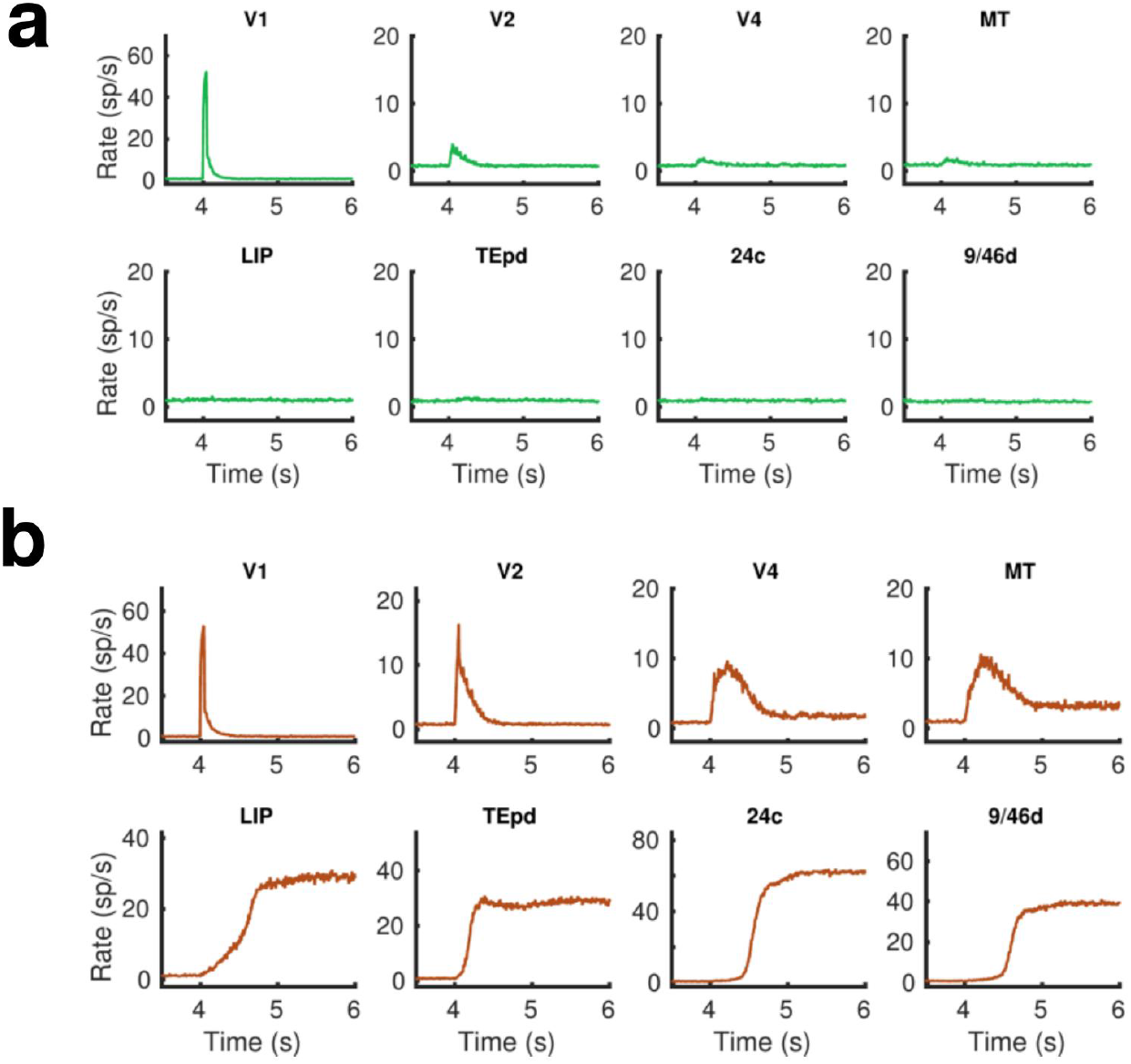
Firing rates for selected areas during a visual WM task with a short (50ms) stimulus duration. (a) Only NMDA and GABA synapses are considered in the network, the stimulus does not reach frontal areas. (b) By introducing simplified AMPA-like synapses (proportional to the firing rates) on feedforward excitatory projections, we obtain distributed WM patterns for this type of short inputs. The results suggests that it is convenient to consider AMPA/NMDA asymmetry along the FF/FB projections in the hierarchy for brief stimulation patterns. Other parameters as in Fig. 2.

**Extended Data Fig. 6:**
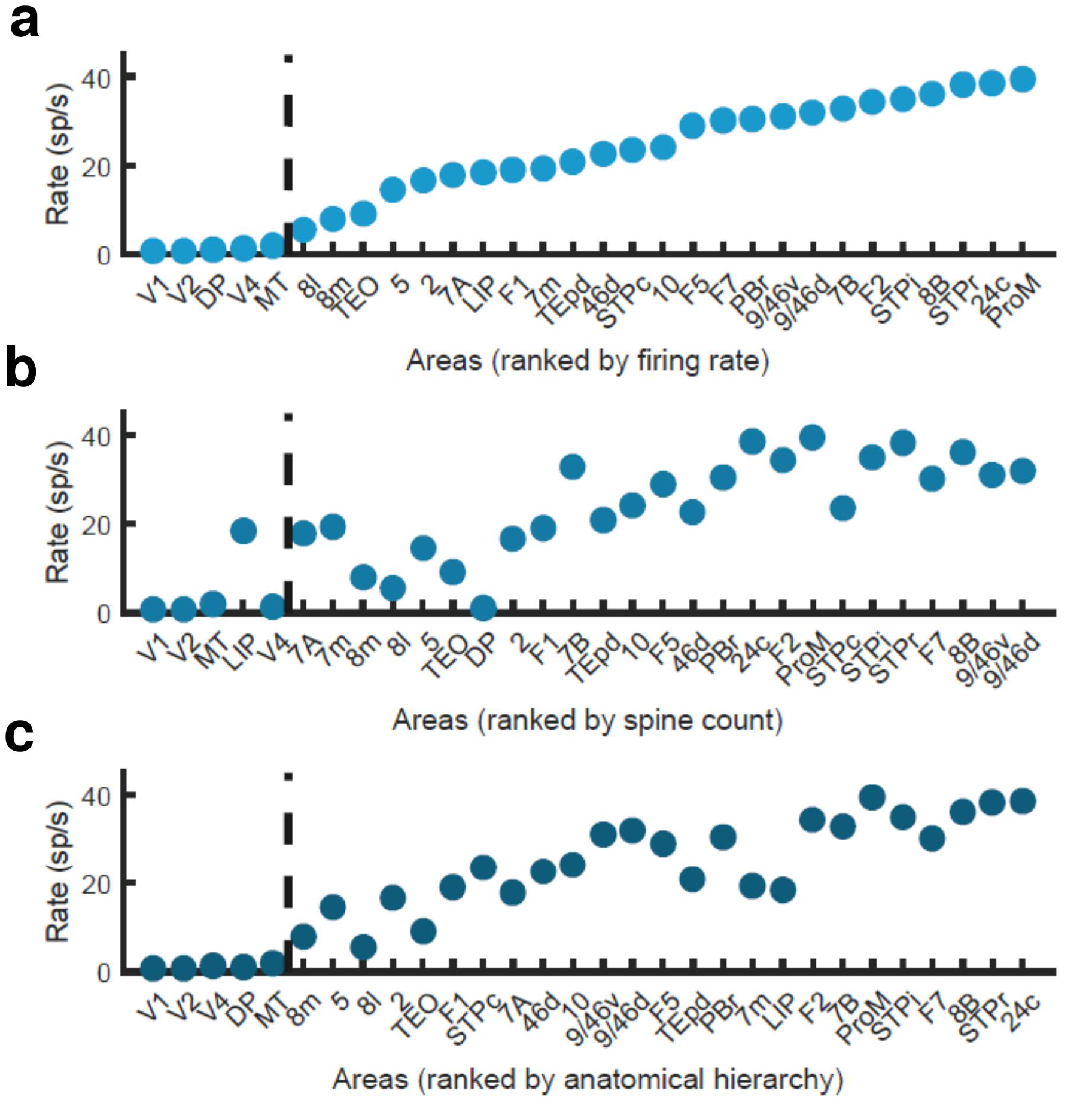
Firing rate of ranked cortical areas reveals a robust bifurcation in space with different ranking systems. (a) Areas ranked by displayed firing rate. (b) Areas ranked by spine count. (c) Areas ranked following their hierarchical (as obtained from SLN data) position. Parameters as in Fig. 2.

**Extended Data Figure 7:**
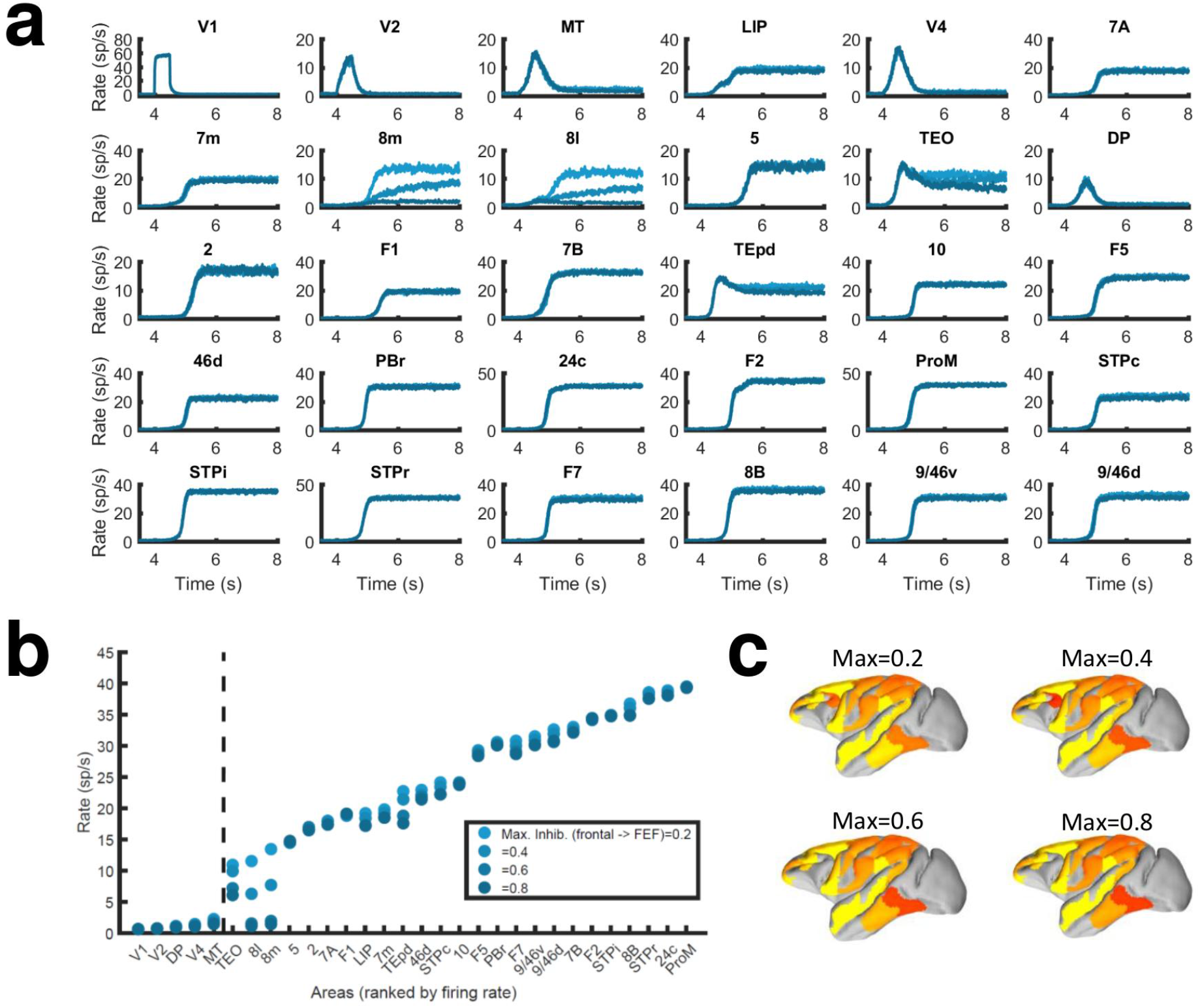
Corrections to localized regions; example of FEF areas. (a) Traces of every area in the simulated neocortical network. We limit here the strength of counterstream inhibitory bias, or preference of feedback towards inhibition, which feedback projections from frontal areas to FEF areas (8l and 8m) may display. For each plotted area, blue lines of increasing darkness show the effects of limiting frontal inhibitory feedback to FEF to 0.2, 0.4, 0.6 or 0.8. The effects are mostly visible in areas 8l and 8m, and to a lesser extent in TEO. (b) Bifurcation in the cortical space for the different limitations to FEF inhibitory feedback. (c) Activity maps for the four conditions shown in (a, b). Other parameters as in Fig. 2.

**Extended Data Fig. 8:**
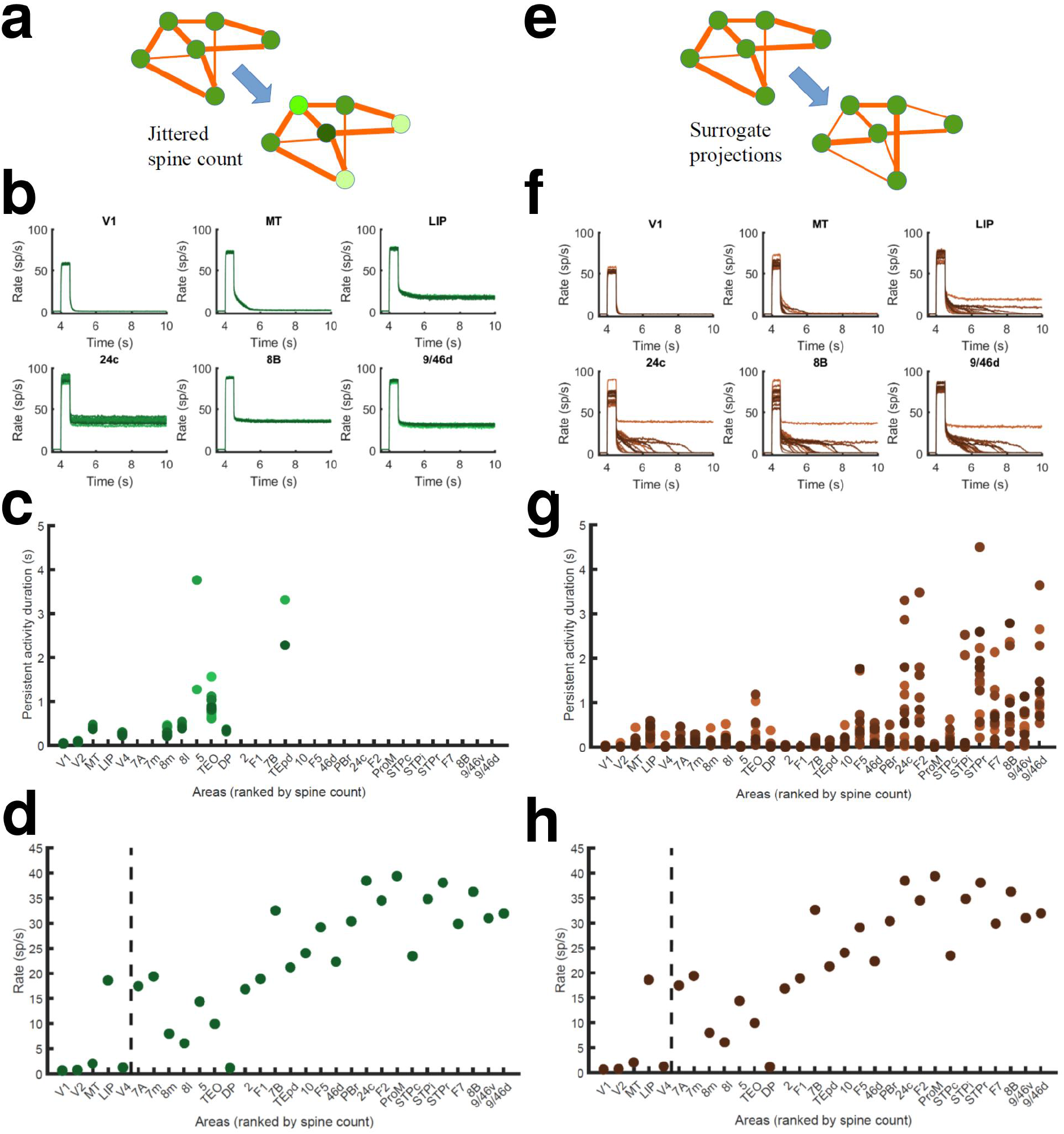
Effect of jittered dendritic spine values and surrogate networks. (a) Scheme of networks with jittered area-specific dendritic spine numbers (within 15% of their original values), and therefore jittered local and long-range synaptic strengths. (b) Effects of the jitter in traces for selected areas (lines of different colors represent activity for different configurations). (c) Jittering has a minor effect on the duration of the persistent state for low/middle spine count areas. (d) Bifurcation in cortical space for jittered networks. (e) Scheme of surrogate networks, in which FLN values are randomly shuffled. (f) Effects of shuffling FLN values on traces of selected areas. (g) Surrogate networks present moderate differences in the duration of the persistent states of high spine count values. (h) Bifurcation in cortical space for surrogate networks. Parameters as in Fig. 2.

**Extended Data Fig. 9:**
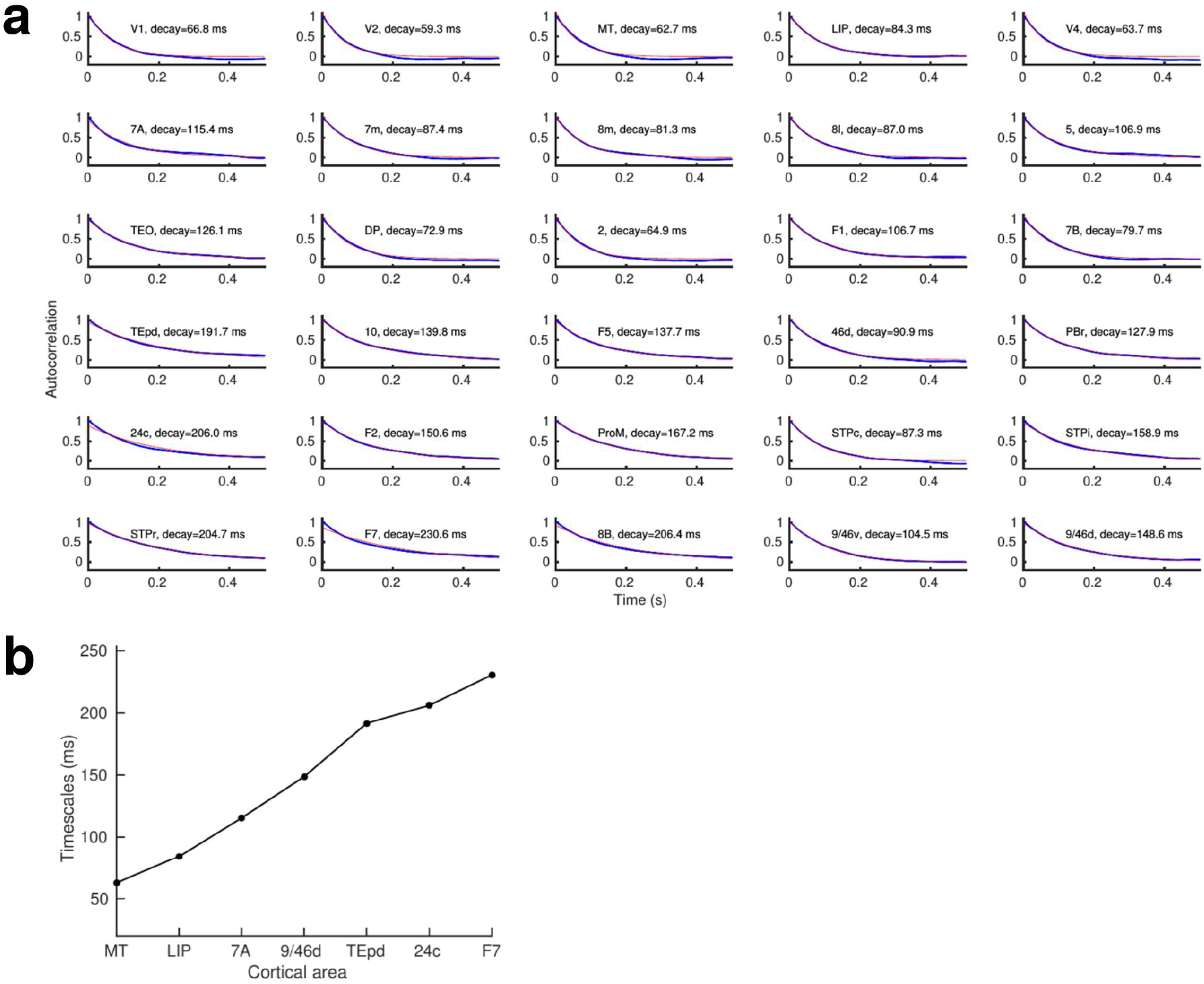
A gradient of time scales emerges in the network. (a) Autocorrelation function of the activity (low-pass-filtered to simulate an LFP recording) of each area in the network, fitted with a decaying exponential function. (b) The values for the local decay time constant increase with the cortical hierarchy, going from ∼60 ms in early visual areas to ∼300 ms in some frontal areas. This range is similar to the spike-count autocorrelations found experimentally across different cortical areas^**31,58**^, and improves predictions from previous models^**21**^ (but see additional considerations^**59**^). Parameters as in Fig. 2.

**Extended Data Table 1:**
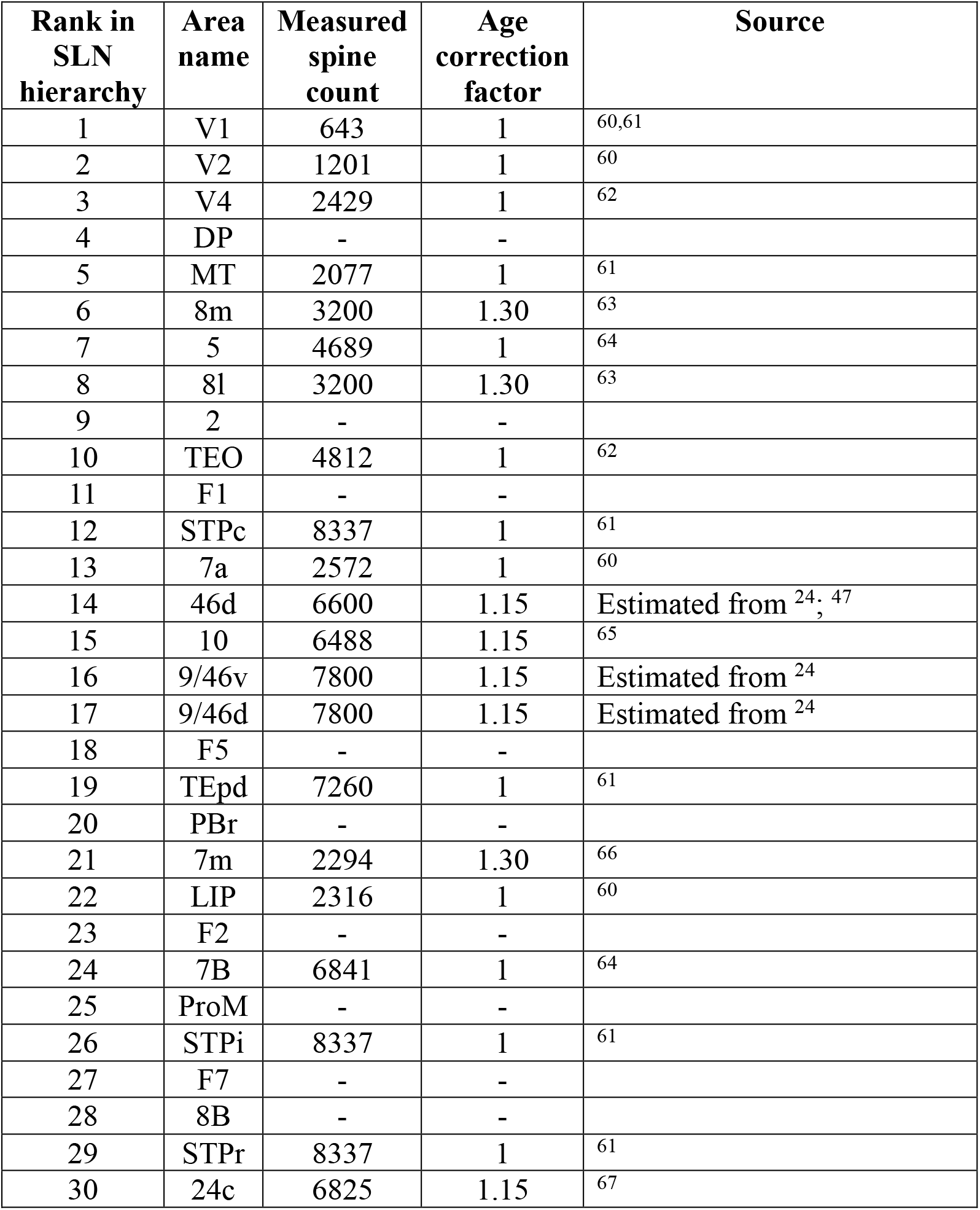
Spine count data from basal dendrites of layer 2/3 pyramidal neurons in young (∼2 y. o.) macaque, acquired from the specified literature. See also Extended Data Fig. 2.

## Notes

### Competing Interest Statement

The authors have declared no competing interest.

